# Shared genomic architecture of the brain white matter structural connectome and intelligence

**DOI:** 10.1101/2025.04.09.647917

**Authors:** Xinyi Dong, Weijie Huang, Haojie Chen, Yunhao Zhang, Bing Liu, Ni Shu

## Abstract

White matter (WM) connections, which facilitate communication between brain regions, substantially impact intellectual performance. However, the shared genetic underpinnings of WM connections and intelligence remain unclear. In the present study, we first conducted genome-wide association studies on the global and regional topological properties of WM connectomes constructed from diffusion-weighted and T1-weighted magnetic resonance images of 26,655 participants from the UK Biobank. Forty-one independent significant single-nucleotide polymorphisms (SNPs) associated with global WM connectome efficiency and 45 SNPs linked to nodal efficiency for 246 brain regions were identified. Then, regional heterogeneity was demonstrated for the genetic correlations between WM connectome efficiency and intelligence. The nodal efficiency of 128 regions with nominally significant genetic correlations with intelligence was determined. Among these, 44 regions and 3 regions shared SNPs with intelligence within the chromosome 6q21 and 3p21.1 locus respectively, and significant causal effects on intelligence were found for 63 regions, mainly in the orbital gyrus and superior frontal gyrus. Finally, we found that the integration of polygenetic score and WM connectome efficiency data was more accurate for predicting individual intelligence than the use of a single data type. These findings improve our understanding of how genes shape the individual WM connectome and intelligence and the relationship between WM connectomes and intelligence.

## Introduction

White matter (WM) connections play critical roles in information transmission between brain regions. The interindividual variability in WM connectomes serves as the neural basis of individual identity, contributing to differences in behavior, cognition, and self-expression^1^. Recently, the human brain connectome has provided an advanced framework for constructing WM structural connectomes from diffusion-weighted magnetic resonance imaging (dMRI) data, enabling studies of the organization of WM connections at the global system level^2^. Specifically, the WM structural connectome has nontrivial topological properties, including small-world, modular and rich-club organization^3–5^. Graph metrics, such as network efficiency^6^, clustering coefficients^7^, and the shortest path length^8^, have been widely applied to quantify the information integration and segregation properties of the brain connectome, and such metrics can elucidate the neural mechanisms underlying individual differences in cognitive functions.

Owing to the increasing availability of large-scale neuroimaging and whole-genome data, neuroimaging genomics has emerged as a rapidly developing scientific field. Through recent large-scale genome-wide association studies (GWASs), a diverse set of genomic loci associated with imaging-derived phenotypes, including regional brain volume^9^, cortical morphology^10,11^, functional connectivity^12^, and WM microstructure^13^, have been identified on the basis of MRI data. Notably, two studies have demonstrated that WM connectivity is heritable to some extent and have provided insights into its genetic architecture^14,15^. The results of both studies suggested that the genes regulating WM connectivity are partially associated with myelination and early brain development. Although the topological properties of WM connectome are derived from WM connectivity, they are not identical to connectivity itself. Instead, these properties can be used to quantitatively characterize the topological organization of WM connectomes. Furthermore, these properties may reflect a trade-off between competing principles: minimizing wiring costs while maximizing adaptive efficiency^16^. Compared with WM connectivity, the topological properties of WM connectomes are less influenced by methodological choices in network construction, such as the selection of parcellation templates^17^. However, the heritability of WM connectome topological properties and its genetic architecture remain largely unknown. Investigating the genetic basis of WM connectome topological properties could provide new insights into how genetic factors shape the relationships between brain connectome and cognition.

Among the various types of human behaviors and cognitive functions, intelligence stands out as a distinct and fundamental construct. Intelligence is one of the most stable cognitive abilities across the lifespan^18^ and is one of the strongest predictors of educational achievement^19,20^, occupational status^21^, mental and physical health^22^, and longevity^23^. The etiology of individual differences in human intelligence has long been a focus in the field of cognitive neuroscience^24^. On the basis of structural and functional MRI studies, several network theories of intelligence have been proposed, such as the parieto-frontal integration theory (P-FIT)^25^ and the multiple demand network^26^. Both theories suggest that intelligence arises from the interaction and collaboration of specific brain regions, including the dorsomedial and lateral prefrontal cortex, insula, and parietal cortex^25,26^. Previous studies have focused primarily on the relationship between structural^27^ and functional changes^28,29^ in these brain regions and individual differences in intelligence while overlooking the influence of WM connectivity among brain regions - the anatomical foundation for interregional information transmission - on intelligence. Additionally, similar to WM connectivity, intelligence is highly heritable and polygenic. Three recent large-scale studies identified 187^30^, 148^31^, and 205^32^ single-nucleotide polymorphisms (SNPs) associated with intelligence, with substantial overlap among these SNPs. Therefore, clarifying the shared genetic architecture between WM connectome topological properties and intelligence is crucial for understanding their relationship and the neural mechanisms underlying individual differences in intelligence.

In this study, we first investigated the genetic architecture underlying WM connectome topological properties. Next, we examined the phenotypic and genetic relationships between WM connectome topological properties and intelligence from multiple perspectives. Finally, we developed models linking genes, WM connectome topological properties, and intelligence, aiming to improve our understanding of their interrelationships. Fig. 1 provides an overview of the study design and analyses.

**Fig. 1.**
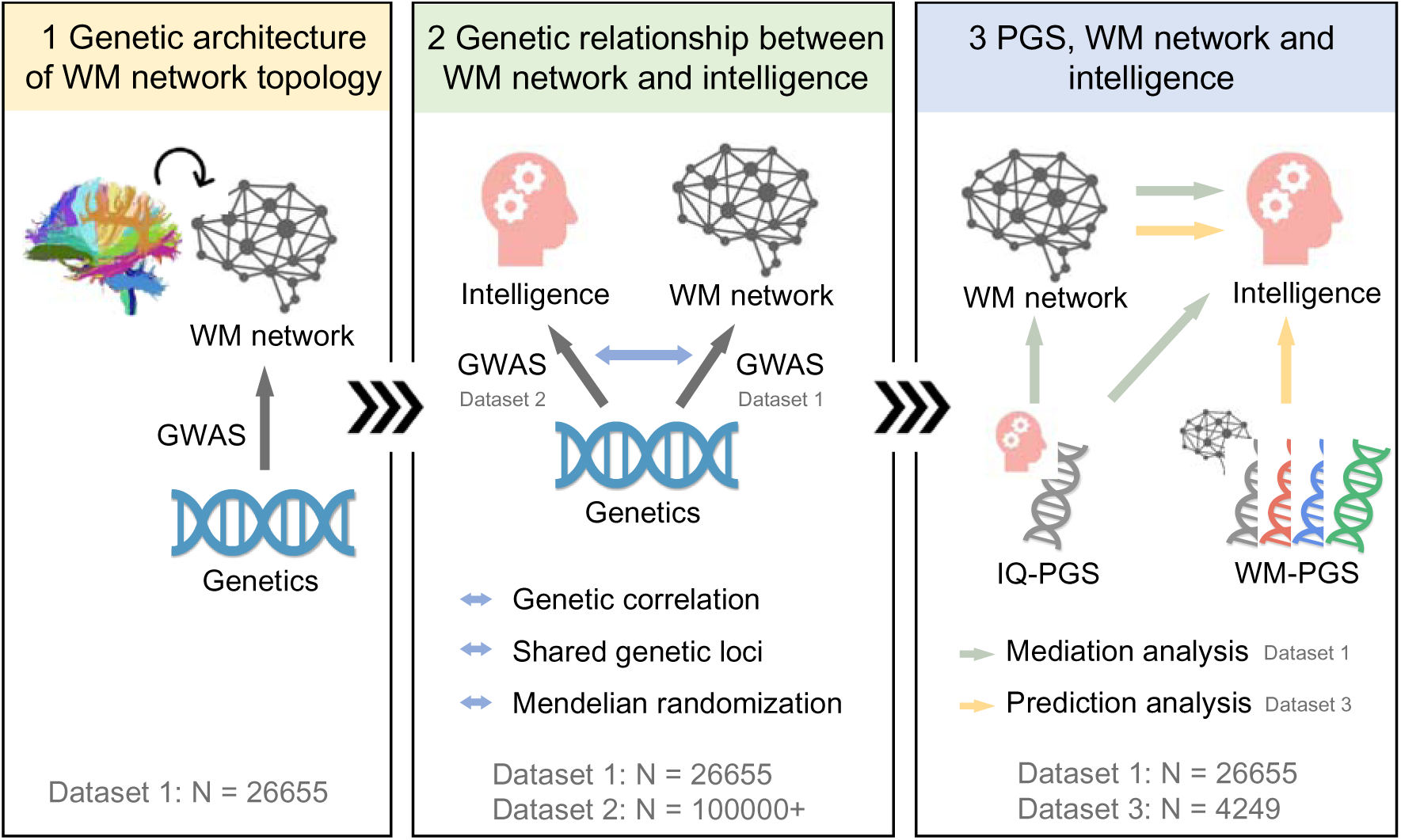
Overview of the study design and analyses. First, we conducted GWAS to explore the genetic architecture underlying WM connectome topological properties. Next, we employed genetic correlation, colocalization and Mendelian randomization to elucidate the genetic relationships between WM connectome topological properties and intelligence. Finally, we calculated the PGSs for intelligence and nodal efficiency and developed mediation and prediction models to explain the interrelationship between genes, WM connectome toplogical properties and intelligence. WM, white matter; GWAS, genome-wide association study; PGS, polygenic score.

## Results

### Genetic architecture of WM connectome topological properties

A total of 26,655 unrelated participants of white British ancestry (aged 46-82 years) from the UK Biobank (UKB)^33^ were included in the GWAS of WM connectome topological properties. For each participant, deterministic fiber tractography was applied to the dMRI data to reconstruct WM tracts; then, a WM connectome (246 X 246 symmetric matrix) was constructed, with using the Brainnetome Atlas (BNA)^34^ to define the network nodes and the number of interconnecting streamlines between regions to define the network edges (Fig. 2a). On the basis of the constructed WM connectome, topological properties were analyzed from both global and nodal perspectives. The global properties, including the global efficiency

**Fig. 2.**
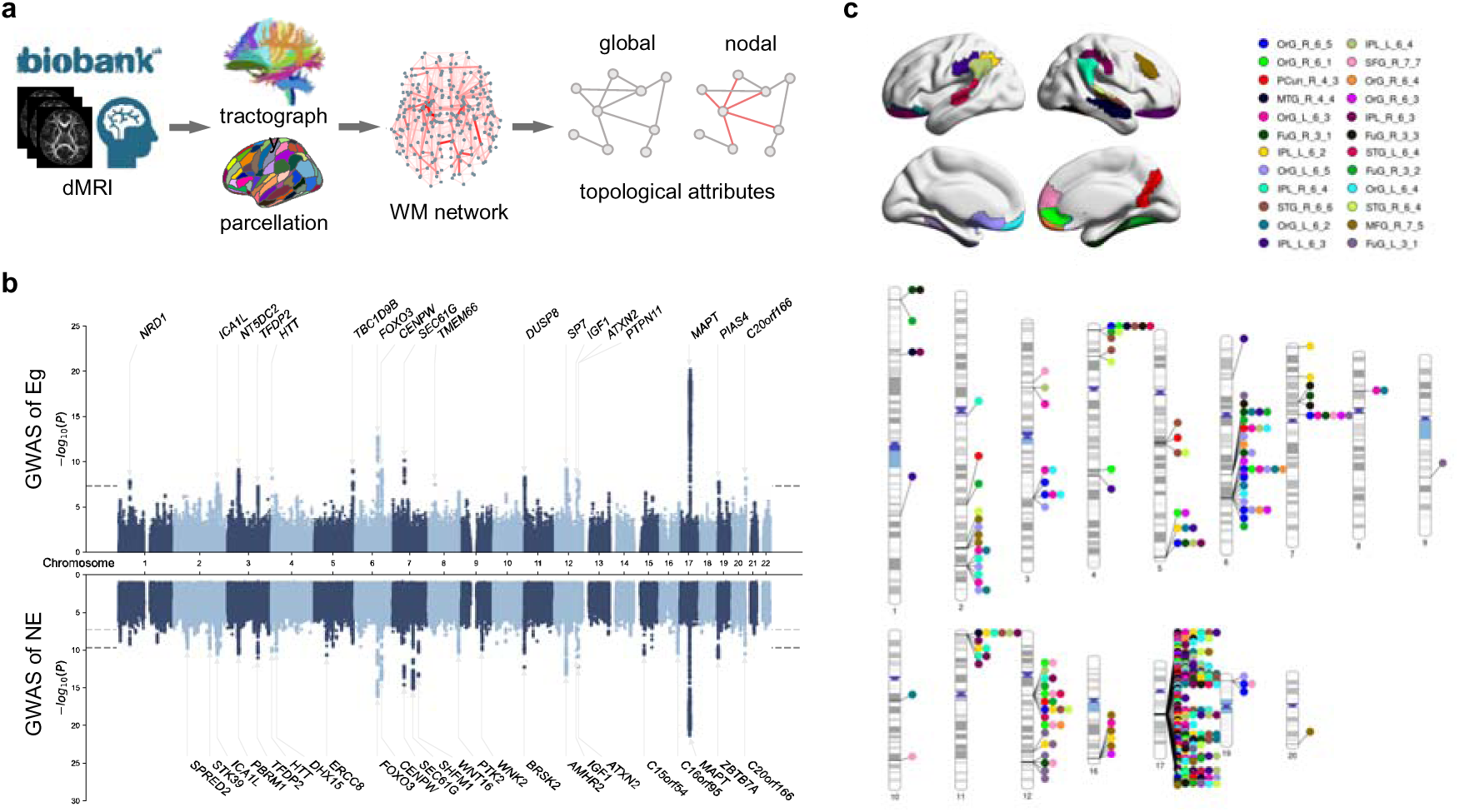
Genetic architecture of WM connectome efficiency. (**a**) Schematic of WM connectome construction and topology. (**b**) Manhattan plots with genetic variants identified through univariate GWAS of global efficiency (upper) and multiple univariate GWAS of NE across 246 regions with min-P approach (lower). The grey lines indicate genome-wide significance threshold (*P* < 5 × 10^−8^), and Bonferroni-corrected threshold for the multiple univariate GWAS (*P* < 2 × 10^−10^). The independent significant variants are annotated by their nearest genes. (**c**) Ideogram of genomic regions influencing NE (*P* < 5 × 10^−8^). The colors represent the 24 regions with the top 10% highest heritability. WM, white matter; GWAS, genome-wised association study; Eg, global efficiency; NE, nodal efficiency; OrG, orbital gyrus; PCun, precuneus; MTG, middle temporal gyrus; FuG, fusiform gyrus; IPL, inferior parietal lobule; STG, superior temporal gyrus; SFG, superior frontal gyrus; MFG, middle frontal gyrus.

(Eg), local efficiency (Eloc), clustering coefficient (Cp), shortest path length (Lp), and small-world parameters (γ, λ, and σ), characterize the overall topological structure of the network. The nodal properties, including the nodal efficiency (NE), nodal local efficiency (NLE), nodal clustering coefficient (NCP), and nodal shortest path length (NLP), describe the roles that each region plays within the network.

We estimated the SNP heritability of the WM connectome topological properties, which represents the proportion of phenotypic variance explained by all genome-wide SNPs. Among all the global properties, Eg presented the highest heritability of 0.36, whereas Cp presented the lowest heritability of 0.10 (Supplementary Fig. 1a). Among the four nodal properties, most brain regions exhibited significant heritability, with NE demonstrating the highest heritability across nearly the entire brain (Supplementary Fig. 1b). The statistical details are provided in Supplementary Table 1.

We performed a univariate GWAS for each global and nodal network properties using 7,301,482 SNPs after genetic quality control. Through the univariate GWAS, we identified 41 independent significant SNPs associated with the Eg of the WM connectome at a significance level of *P* < 5 × 10^-8^ (Fig. 2b, Supplementary Table 2). The genetic architectures of Eloc and Lp were similar to that of Eg, although the SNP effects were weaker for these properties (Supplementary Figs. 2 and 3). In contrast, different genomic loci were identified for the Cp and small-world parameters (Supplementary Figs. 4 and 5). For the nodal properties, we identified 45 independent significant SNPs associated with at least one of the NE of the 246 brain regions after Bonferroni correction (*P* < 5 × 10^−8^/246 = 2 × 10^-10^, Fig. 2b, Supplementary Table 3). These SNPs were linked to 24 genes, 10 of which were also identified in the GWAS of Eg. Additionally, 222 independent significant SNPs were identified at a nominal genome-wide significance threshold (*P* < 5 × 10^−8^, Fig. 2b, Supplementary Table 3). From the GWAS results of the NE across the 246 brain regions, we highlighted the genomic regions associated with the brain regions with the top 10% highest heritability, each reaching genome-wide significance (*P* < 5 × 10^-8^, Fig. 2c). These regions included the orbital gyrus (OrG), precuneus (Pcun), middle temporal gyrus, fusiform gyrus, inferior parietal lobule (IPL), superior temporal gyrus (STG), superior frontal gyrus (SFG), and middle frontal gyrus (MFG). The associated genomic regions were predominantly located on chromosome 17. Furthermore, the genetic architectures of the NLE and NLP closely resembled that of NE, whereas a distinct genetic architecture was found for the NCP, with its genetic loci located primarily on chromosome 7 (Supplementary Figs. 2-4).

We performed gene-based association analyses using the minimum p value from the GWASs of the nodal properties across the 246 brain regions with MAGMA^35^ implemented in FUMA^36^ (Supplementary Table 4-7). For NE, 7,455 significant genes were identified (P < 0.05/19279/246 = 1.0 × 10^-8^; Supplementary Table 4). The significant genes identified by MAGMA were investigated further through enrichment analysis. The gene ontology (GO) gene sets obtained from MsigDB^37^ revealed that the genes were significantly enriched for 159 biological process terms, 27 cellular component terms, and 26 molecular function terms (Benjamini[Hochberg adjusted P < 0.05, Supplementary Table 8). The top enriched terms, shown in Supplementary Fig. 6, were related primarily to signal transduction, neuron projection, and ion transport. The enriched terms for the other nodal properties were largely similar to those identified in the NE enrichment analysis (Supplementary Tables 9-11). Additionally, we conducted a drug target enrichment analysis and that drug targets for nervous system diseases, eye diseases, and mental, behavioral, neurodevelopmental disorders were significantly enriched (ICD-10 codes; Supplementary Fig. 7) and cardiovascular disease drugs (ATC codes; Supplementary Fig. 8).

### Phenotypic and genetic correlations between WM connectome topological properties and intelligence

We next examined the phenotypic and genetic associations between WM connectome topological properties and intelligence in the participants from the UKB. Pairwise phenotypic associations in unrelated white British individuals who underwent both dMRI scans and fluid intelligence tests (n = 24,704) were investigated using partial correlation analysis, controlling for sex, age and assessment center. Among the global properties, Eg appeared to be the most strongly related to intelligence (r_p_ = 0.12, P = 1.24 × 10^-83^, Supplementary Table 12-1). Regional correlation analyses revealed a significant association between NE and intelligence across all brain regions (FDR-corrected P < 0.05, Fig. 3a). This correlation was particularly strong in the SFG, OrG, IPL, temporal lobe, and subcortical regions, including the basal ganglia and thalamus, closely aligning with the correlation patterns observed for the NLP and NLE (Supplementary Fig. 9). However, weaker correlations between the NCP and intelligence were observed across the whole brain (Supplementary Fig. 9). To investigate the genetic correlations between WM connectome topological properties and intelligence, we conducted a GWAS of intelligence in 182,798 unrelated white British individuals. Through the GWAS, we identified 301 independent significant SNPs associated with intelligence at a genome-wide significance level of P < 5 × 10^-8^ (Fig. 3b, Supplementary Table 13). The associated genomic regions were predominantly located on chromosome 6, with one of the top signals associated with the FOXO3 gene, which was also identified in the GWAS of the Eg of the WM connectome. We then estimated genetic correlations between WM connectome topological properties and intelligence using linkage disequilibrium score regression (LDSC). Consistent with the phenotypic correlations, Eg and Eloc presented the strongest genetic associations with intelligence (Supplementary Table 12-2). Regional LDSC analyses further revealed significant genetic correlations between various nodal properties and intelligence (Supplementary Fig. 10). Specifically, the NE in 128 brain regions was nominally significantly associated with intelligence (Supplementary Fig. 11), and these associations remained significant in 93 of the 128 regions following FDR correction (Fig. 3c), including the SFG, OrG, IPL, Pcun, paracentral lobule (PCL), medioventral occipital cortex (MVOcC), left insular gyrus (INS), and subcortical regions, such as the basal ganglia and thalamus. Considering that network efficiency has the highest heritability and the strongest phenotypic and genetic correlation with intelligence, we focused on Eg and NE to explore the genetic mechanisms shared between WM connectome topological properties and intelligence in subsequent analyses.

**Fig. 3.**
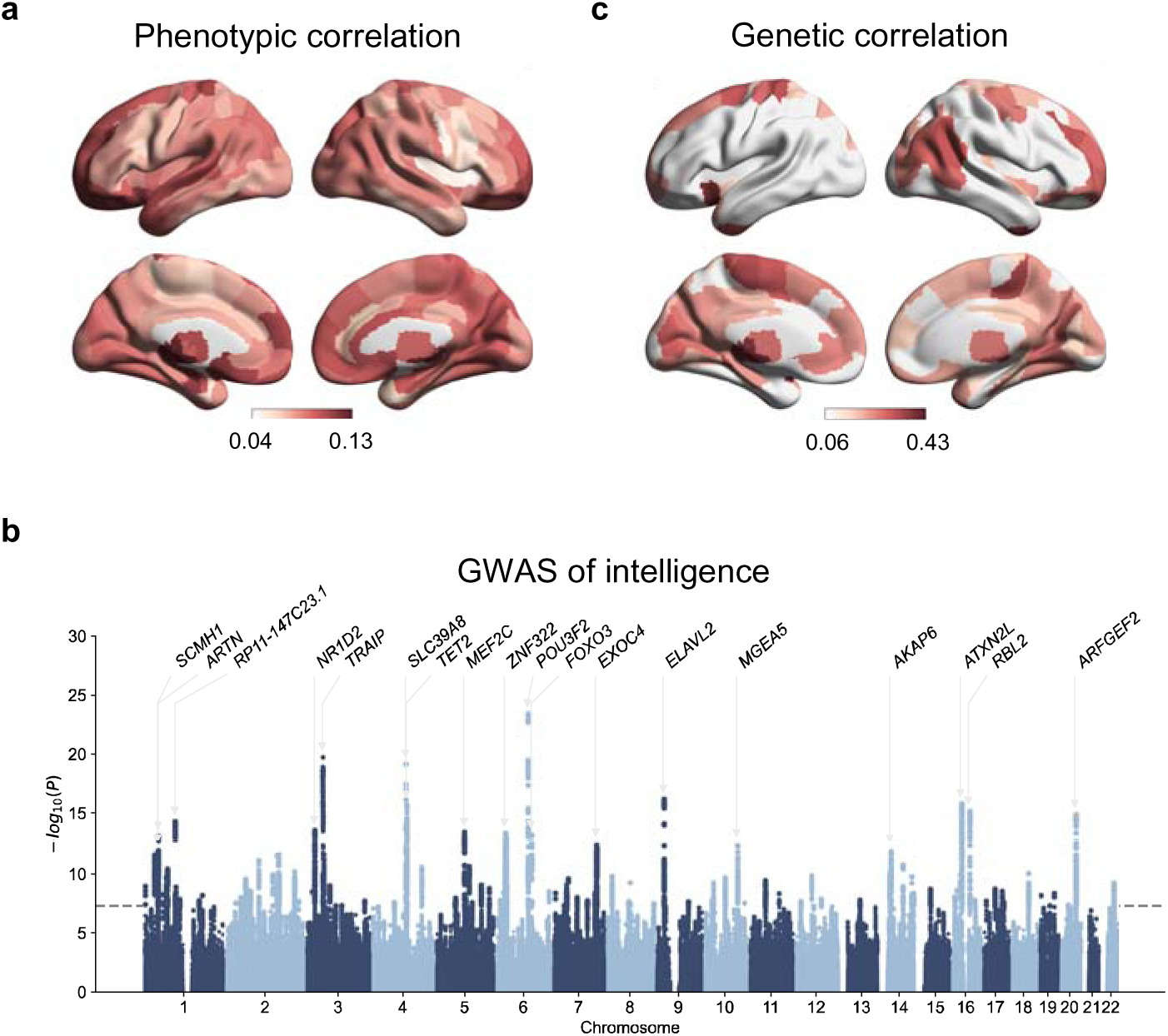
Phenotypic and genetic correlations between WM connectome efficiency and intelligence. (**a**) The phenotypic correlations between nodal efficiency and intelligence were estimated controlling for sex, age and assessment center. (**b**) Manhattan plots with genetic variants identified through univariate GWAS of intelligence. The grey line indicates genome-wide significance threshold (P < 5 × 10^−8^). The independent significant variants are annotated by their nearest genes. (**c**) The genetic correlations between nodal efficiency and intelligence were estimated using LDSC. Regions with significant genetic correlations with intelligence (false discovery rate corrected P < 0.05) are highlighted. WM, white matter; GWAS, genome-wised association study; LDSC, LD-score regression.

### Shared genetic loci between WM connectome efficiency and intelligence

To evaluate whether WM connectome efficiency and intelligence share causal variants, we conducted a Bayesian colocalization analysis^38^. Colocalization was defined by a posterior probability greater than 0.75, indicating a shared causal variant. Among the forty-one independent SNPs associated with Eg, seven SNPs were shared with intelligence, spanning chromosomes 3, 6, and 17 (Supplementary Table 14). For the 128 regions that demonstrated nominally significant genetic correlations with intelligence, colocalization analyses were performed to determine whether these regions share causal variants with intelligence. Similar to Eg, most loci were shared between NE and intelligence resided on chromosomes 3, 6, and 17 (Supplementary Table 15). Additionally, three SNPs on chromosome 2 and one SNP on chromosome 5 were shared between NE and intelligence. The regions shared SNPs that correlated with NE with *P* values < 5 × 10^-8^ are shown in Fig. 4a. Among these, 44 SNPs on chromosome 6q21 were shared with intelligence, mapping to regions including the lateral SFG (Fig. 4c), left OrG (Fig. 4d), caudal dorsolateral precentral gyrus, right ventral fusiform gyrus (FuG), trunk region in the postcentral gyrus, MVOcC, lateral occipital cortex, basal ganglia and thalamus. Additionally, three brain regions shared SNPs on chromosome 3p21.1 with intelligence, including the left dorsal area of the MFG, left medioventral area of the FuG, and left caudal hippocampus (Fig. 4b). On chromosome 6q21, colocalization was identified in the FOXO3 gene, where independent significant variants of this gene have been associated with intelligence, educational attainment, schizophrenia, body mass index, brainstem volume, and lung function, as reported in the National Human Genome Research Institute–European Bioinformatics Institute GWAS Catalog^39^.

**Fig. 4.**
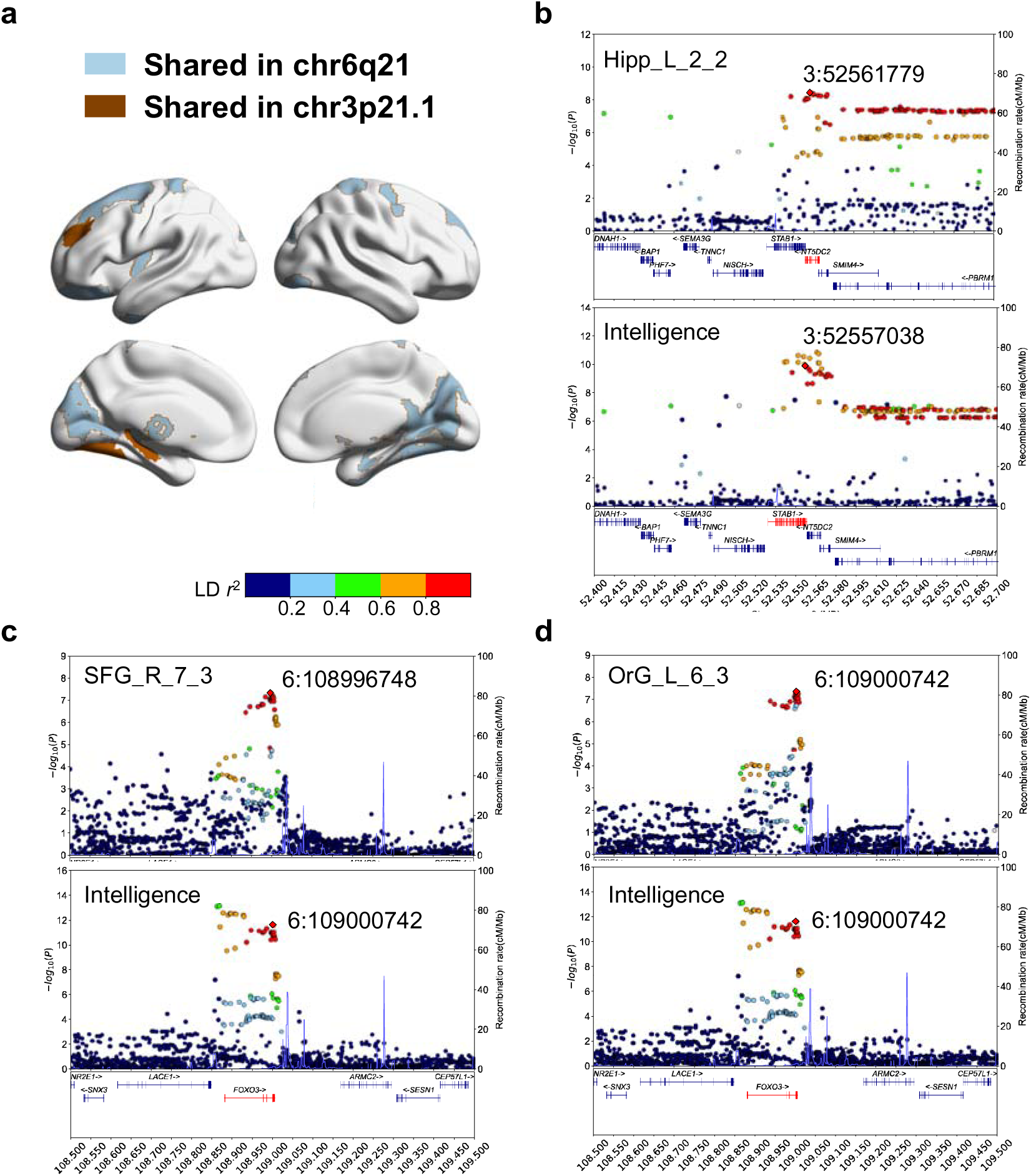
Genetic loci associated with nodal efficiency of WM connectome and intelligence. (**a**) The regions sharing variants with intelligence are highlighted. The two colors represent colocalizations occurring at different loci. (**b**) In chr3p21.1, we observed colocalization between the efficiency of left caudal Hipp (chr3:52561779) and intelligence (chr3:52557038). (**c**) In chr6q21, we observed colocalization between the efficiency of right lateral SFG (chr6:108996748) and intelligence (chr6:109000742). (**d**) in chr6q21, we observed colocalization at variant chr6:109000742 between the efficiency of left lateral OrG and intelligence. WM, white matter; Hipp, hippocampus; SFG, superior frontal gyrus; OrG, orbital gyrus.

### Causal relationships between WM connectome efficiency and intelligence

To further investigate the causal relationship between WM connectome efficiency and intelligence, we employed a two-sample Mendelian randomization (MR) approach. At the global level, the causal effects of Eg on intelligence were significant across all MR methods after Bonferroni correction (P < 2.8 × 10-3, correcting for 9 MR methods and 2 causal directions; Fig. 5c, Supplementary Table 16). In contrast, the reverse effects of intelligence on Eg were not significant in any MR methods (Supplementary Table 16). At the regional level, for the 128 regions showing nominally significantly genetic correlation with intelligence, we conducted MR analyses to assess the causal relationship between the NE and intelligence. Forward MR analysis identified 63 regions with causal effects on intelligence, including OrG, right STG, right IPL, left INS, MVOcC, MFG, basal ganglia and thalamus (Fig. 5d, Supplementary Table 17). In contrast, reverse MR analysis did not identify any significant causal effects of intelligence on NE, further supporting the directional influence of WM connectome efficiency on intelligence.

**Fig. 5.**
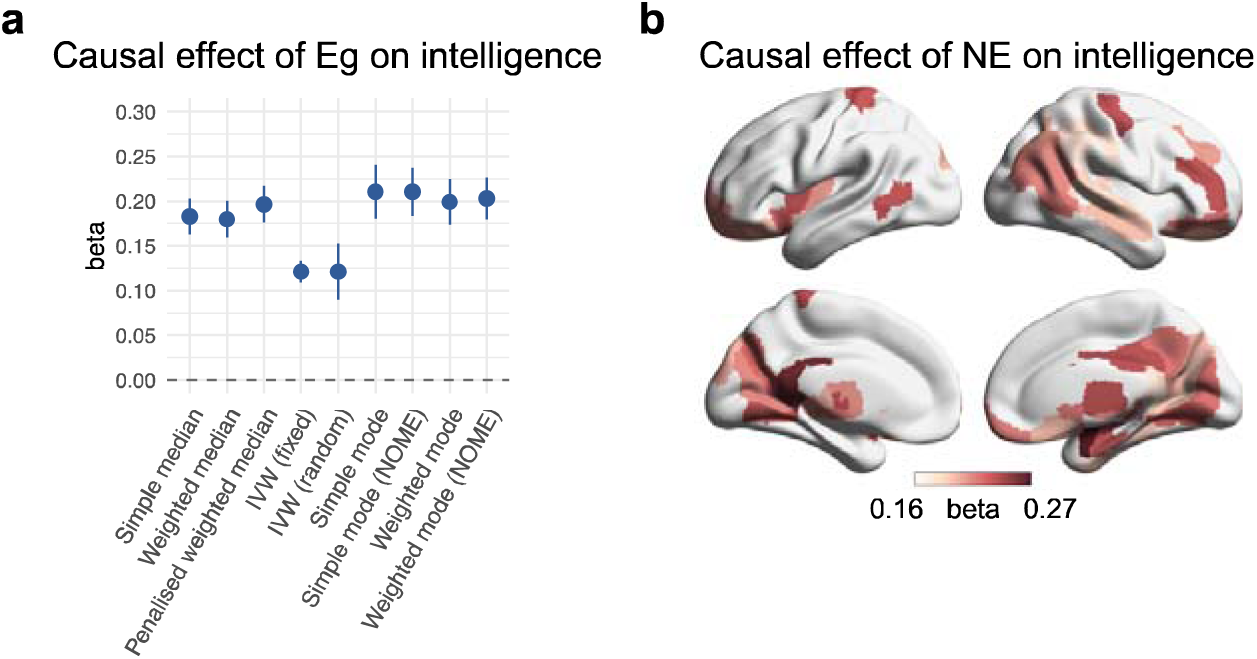
Causal effects of WM connectome efficiency on intelligence. **(a)** The significant causal relationships from Eg (exposure) to intelligence (outcome) using nine MR methods after Bonferroni correction (*P* < 2.8 × 10^-3^). (**b**) The regions where causal relationships from NE (exposure) to intelligence (outcome) were significant after Bonferroni correction (*P* < 1.1 × 10^-5^) were colored according to the beta value, representing the causal effect. WM, white matter; INT, intelligence; PGS, polygenic score; Eg, global efficiency; NE, nodal efficiency.

To assess the robustness and reliability of the forward MR results, we conducted sensitivity analyses for the significant MR findings. First, a leave-one-out analysis for Eg and NE revealed that no single SNP drove the observed associations (FDR-corrected P < 0.05; Supplementary Table 18), except for one region in the basal ganglia, where the MR result was already marginally significant. Second, MR-Egger intercepts of the associations were close to zero for both Eg and NE, indicating no significant horizontal pleiotropy (FDR-corrected P > 0.05; Supplementary Table 19). This confirms that the exclusivity and independence assumptions of MR analysis were satisfied. Third, the MR results for Eg failed the heterogeneity tests (P < 0.05; Supplementary Table 20), and MR estimates from individual SNPs showed both positive and negative effects, suggesting inconsistent SNP-level effects (Supplementary Fig. 12, Supplementary Table 21). At the regional level, 34 out of 63 regions failed the heterogeneity tests (FDR-corrected P < 0.05; Supplementary Table 20). Despite this heterogeneity, the overall sensitivity analyses confirm the reliability and robustness of our forward MR results.

### Mediation effect of WM connectome efficiency on the relationship between polygenic score and intelligence

Based on the GWAS result of intelligence, the polygenic score (PGS) reflecting polygenic effects on intelligence was computed in an independent sample of white British individuals (N = 26,655). The PGS for intelligence (INT-PGS) accounted for 5.11% of the phenotypic variation in intelligence (*P* = 7.41 × 10^−284^). We then explored the mediating role of WM connectome efficiency in the relationship between INT-PGS and intelligence. The mediation effect was considered significant if the 95% confidence interval (CI) of the bootstrap did not include zero. At the global level, mediation analysis showed that Eg significantly mediated the relationship between INT-PGS and intelligence (ab[=[0.006, 95% CI[=[[0.004 to 0.007], Fig. 6a). At the regional level, we conducted regional mediation analyses in 128 brain regions that exhibited significant genetic correlations between NE and intelligence to identify specific brain regions involved in the relationship. Among these, 113 regions demonstrated significant mediation effects, accounting for more than 0.8% of the total effect (Fig. 6b). These regions included the SFG, OrG, right IPL, left INS, PCun, MVOcC, MFG, FuG, basal ganglia and thalamus.

**Fig. 6.**
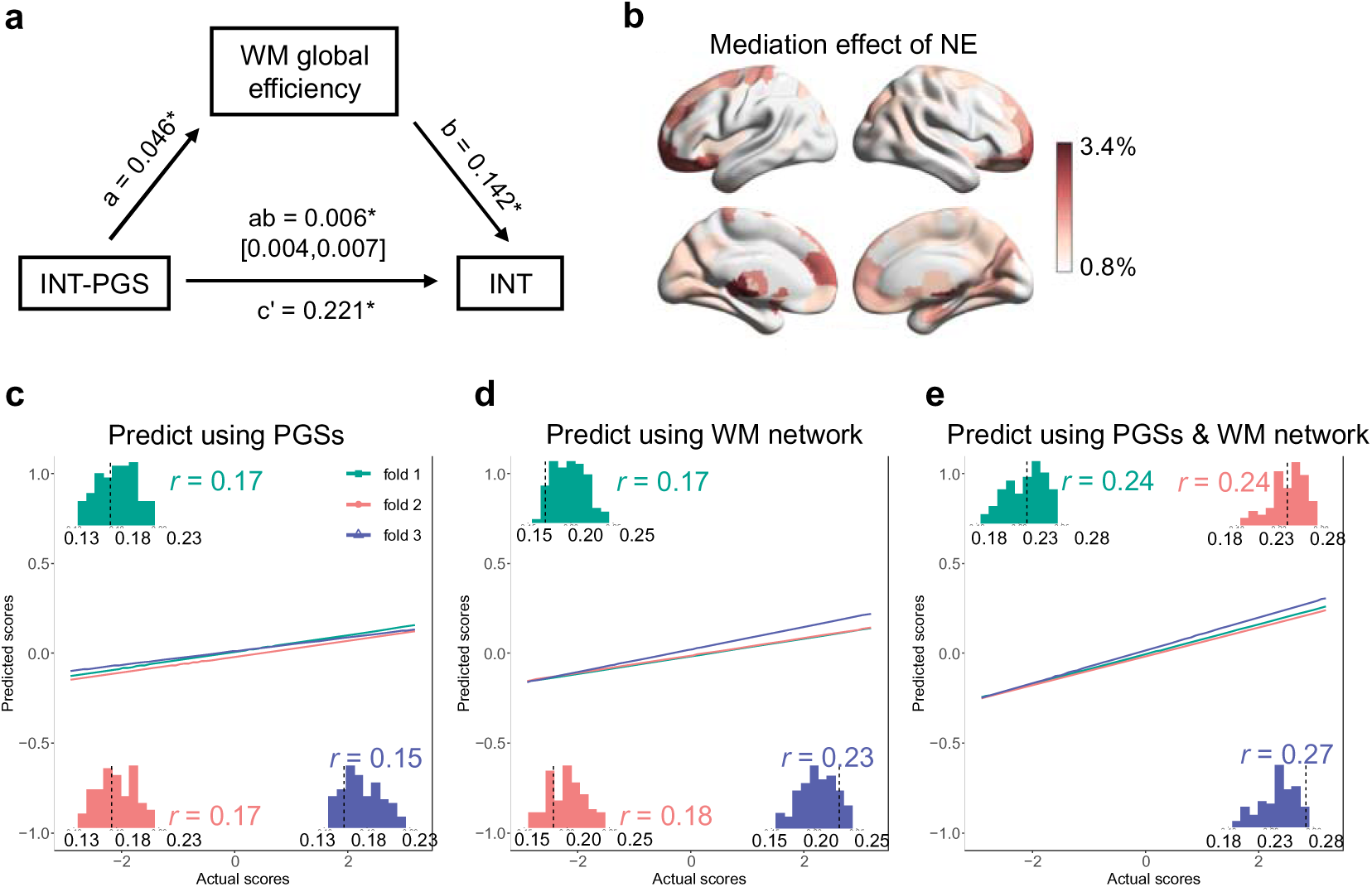
The interrelationship between PGS, WM connectome efficiency and intelligence. (**a**) The mediating role of the global efficiency on the relationship between INT-PGS and intelligence. (**b**) Regions where the nodal efficiency significantly mediates the relationship between INT-PGS and intelligence were colored according to the proportion of the mediating effect. The mediation analyses were performed through 5,000 bootstrapping. An indirect effect was considered significant if the bootstrapped confidence interval did not include zero. The prediction of intelligence using **(c)** PGSs for the nodal efficiency of WM connectome, **(d)** nodal efficiency of the WM connectome, and **(e)** a combination of both as features were shown. The performance of modes were measured by Pearson correlation between predicted and actual intelligence scores. The predictions were obtained through 100 iterations of 3-fold cross-validation using PLS regression. The scatter plots and the correlation values aside depict the results from one cross-validation iteration, with colors representing different folds. The histograms within each fold illustrate the distribution of correlation values between predicted and actual scores across the 100 repetitions. WM, white matter; INT, intelligence; PGS, polygenic score; Eg, global efficiency; NE, nodal efficiency; PLS, partial least squares.

### Prediction of intelligence

Given the phenotypic and genetic associations between WM connectome efficiency and intelligence, we further evaluated the predictive utility of the network efficiency and genetic factors for individual intelligence scores. Using GWAS results for nodal efficiency in white British participants from UKB, we calculated PGS for WM connectome efficiency (WM- PGS) in an independen sample of unrelated individuals from other ancestries (N = 4,249). We adopted partial least squares regression for prediction analysis, using WM-PGSs, NE, and their combination as features, respectivly. The predictive performance was assessed by computing Pearson correlations between predicted and actual intelligence scores, over 100 repeated 3-fold cross-validation. Results showed that using WM-PGSs alone yielded an average correlation of *r* = 0.18 (range: 0.12–0.24, Fig. 6c), while WM connectome efficiency produced *r* = 0.20 (range: 0.14–0.27, Fig. 6d). The model incorporating both yielded an average correlation of *r* = 0.24 (range: 0.14–0.30, Fig. 6e), significantly outperformed models using network efficiency alone (*P* = 1.3 × 10^-48^) and WM-PGSs alone (*P* = 1.7 × 10^-78^).

### Reproducibility analyses

To evaluate the robustness of the genetic architecture of the WM connectome and its genetic correlation with intelligence, we conducted analyses using two independent half-sample subsets of the discovery dataset (dataset 1-1: n = 13327; dataset 1-2: n = 13328). Each of the 41 independent significant SNPs identified in the discovery GWAS of Eg was tested. In dataset1-1, 6 SNPs reached genome-wide significance (*P* < 5 × 10^-8^), while 32 SNPs were significant at the threshold of *P* < 2 × 10^-4^ (0.05/246; Supplementary Table 22, Supplementary Fig. 13). Similarly, in dataset1-2, 8 SNPs reached genome-wide significance (*P* < 5 × 10^-8^), while 32 showed association at the significance level of *P* < 2 × 10^-4^ (Supplementary Table 23, Supplementary Fig. 14). All associations exihibited consistent effect directions with those in the discovery GWAS. LDSC analysis confirmed significant genetic correlations between Eg and intelligence in both half-sample subsets (dataset1-1: r_g_ = 0.20, *P* = 0.02; dataset1-2: r_g_ = 0.27, *P* = 0.01). At the regional level, the number of regions with significant genetic correlations was reduced but remained largely consistent with those in the discovery GWAS (Supplementary Fig. 15-16). These findings suggest that although the reduced sample size of the half-sample datasets limited the statistical power to replicate all significant associations, it was sufficient to replicate a subset of the most prominent results, further supporting the robustness of our conclusions.

We also assessed the robustness of the GWAS results across different brain parcellation templates. Using the Automated Anatomical Labeling atlas that parcellates the cerebrum into 90 regions^40^, we repeated the network construction and GWAS procedures. Among the 41 independent significant SNPs identified in the discovery GWAS of Eg, 26 SNPs remained significant at the level of *P* < 5 × 10^-8^ in the replication GWAS (Supplementary Table 24), and the genetic architecture of the two GWAS was highly consistent (Supplementary Fig. 17). In addition, LDSC analysis revealed a significant genetic correlation between Eg and intelligence (r_g_ = 0.23, *P* = 7.9 × 10^-5^). At the regional level, the spatial distribution of regions showing significant genetic correlations with intelligence was consistent with the findings from BNA atlas, predominantly located in SFG, PCun, PCL, MVOcC, IPL, and OrG (Supplementary Fig. 18). These findings demonstrate the strong generalizability of our GWAS results across different parcellation templates.

## Discussion

In this study, we described the genetic architecture of WM connectome topological properties and revealed the shared genetic mechanism between WM connectome topological properties with intelligence across 26655 UK Biobank participants. We identified 41 independent SNPs associated with Eg, and 45 independent SNPs associated with NE. Among of the 41 independentt SNPs associated with Eg, 7 SNPs were colocalized with SNPs associated with intelligence. Regional analyses showed the heterogeneity in genetic correlation between nodal properties with intelligence across brain regions. Furthermore, we revealed the causal relationship from WM connectome efficiency to intelligence. Finally, we found both WM- PGS and INT-PGS contained the shared genetic component that can predict both intelligence and WM connectome efficiency.

As we hypothesized, WM connectome topological properties showed certain heritability. Particularly, Eg of WM connectomes, which measures the capacity of parallel information transfer in the network and highly relates to cognitive performance, exhibited the highest heritability. This suggested a considerable of genetic influence on the human brain’s ability to process information in parallel. When focusing on NE, the hub areas including Pcun, MTG and IPL were the most heritable, which supports the theory that genes play a preferential role in shaping connections between connectome hubs^41^.

Among the top 10 independent SNPs significantly associated with Eg of WM connectomes, 3 of them (rs62062288, rs9899833 and rs8067056) lies within the MAPT gene, which has been reported to relate to the WM connectivity^15^. The MAPT gene encodes a protein called tau, whose roles is primarily maintaining the stability of microtubules in axons. Aberrant tau protein in neurons has been proved to be associated with several neurodegenerative disorders like Alzheimer’s disease, frontotemporal dementia and cortico-basal degeneration^42^. Notably, rs9899833 is the only SNP that has been reported to associate with WM microstructure^43^, while rs62062288 and rs8067056 are correlated to cortical area^44^ and brain volume^45^ respectively. 2 of them (rs4280294 and rs199443) lies within the NSF gene. The NSF gene encodes a protein that is able to bind to the GluR2 subunit AMPA glutamate receptors, thus it plays a critical role in delivery and expression of AMPA receptors at synapses^46^. This suggests rs4280294 and rs199443 may influence the Eg of WM connectomes by regulating the function of AMPA receptors at synapses. 2 of them (rs450237 and rs12938031) lies within the LINC02210 gene which is an RNA gene. Although its’ function is unclear yet, both SNPs showed significant association with cerebral structure^44,45^. The top fifth SNP, rs3800228, lies within the FOXO3 gene. The FOXO3 gene involves in various functions like infertility, lymphoproliferation, metabolism, organ inflammation etc. FOXO3 signaling pathway regulates the maturation of oligodendrocyte progenitor cells, thus influencing the differentiation of oligodendrocyte progenitor cells and myelination of axon^47^. The top sixth SNP, rs2696694, lies within the LRRC37A gene, which is selectively expressed in pyramidal neurons of the human cerebral cortex and decreases the excitability of the neurons. And rs2696694 was reported to have significant correlation with WM microstructure^48^. Although the top tenth SNP, rs76928645, is an intergenic variant, multiple studies consistently indicated that it is associated with brain voluem^45,49–52^. Altogether, Eg of WM connectomes might be mainly influenced by genes that are (i) related to WM microstructure; (ii) associated with macrostructural characteristics like cerebral volume and cortical area; and (iii) involved in the regulation of neuronal function activity and development.

Gene set analysis also hints roles for neural function and development, which is crucial in the growth, connectivity, signal transmission, and survival of neurons. For example, we found the genes related to NE of WM connectomes were significantly enriched in the GO term ‘regulation of neuron projection development’, a biological process about the development of neurites. This suggests the properties of macroscopic WM connectomes closely link to the development of microscopically neural connections. Other closely related GO terms, such as ‘transmembrane receptor protein tyrosine kinase signaling pathway’, ‘positive regulation of protein kinase b signaling’, ‘positive regulation of intracellular signal transduction’, ‘positive regulation of MAPK cascade’ and ‘cell-cell adhesion mediated by cadherin’, are associated with neural function, including signal transmission, survival and plasticity of neurons. Besides these neuron-specific biological processes, we also identified several biological processes, broadly presenting in various cell types, were associated with the NE of the WM connectomes. Therefore, the biological mechanisms underlying the specific topological structure of the human brain networks are complex and are regulated by multiple biological processes.

We replicated the association between NE and intelligence, which has been reported by several studies^53–55^. The inconsistency between our results and previous findings is that we found significant correlation between NE and intelligence across the whole brain rather than several specific brain regions. This may be due to our large sample size, which allows to detect very subtle associations. By revealing a more global pattern of network efficiency, our study advances the field, demonstrating that intelligence relies on the efficient integration of information across widespread brain regions. The genetic correlation between network efficiency and intelligence suggests the shared genetic mechanism may partly contribute to the phenotypic correlation.

In addition to identifying this genetic correlation, we used colocalization analysis to pinpoint shared genetic loci between network efficiency and intelligence. These loci are primarily located on chromosome 3 (TFDP2, ALAS1 and NT5DC2), 6 (FOXO3 and CENPW) and 17 (LRRC37A). Most of these genes are involved in multiple biological processes, except for ALAS1, which primarily regulates heme biosynthesis. TFDP2^56–58^, FOXO3^59–61^, NT5DC2^62^, and CENPW^63,64^ are mainly involved in regulating cell cycle, proliferation, and apoptosis. Meanwhile, LRRC37A^65^ are potentially involved in biological processes related to the development of the nervous system. The pleiotropy of these genes with respect to WM connectome efficiency and intelligence might be attributed to the pleiotropy of gene products, tissue-specific gene expression, and the complexity of regulatory networks. For example, the products of TFDP2 and FOXO3 are both transcription factors, which can form complexes with other transcription factors to regulate the expression of different target genes, thereby influencing various biological processes. These genes are involved in complex biological processess, thus the biological pathway of these genes regulating network efficiency and intelligence still require further investigation using multi-omics data such as epigenetics and proteomics.

Region-wise analyses further support the idea of shared genetic mechanisms between network efficiency and intelligence by identifying 128 regions nominally genetically associated with intelligence. These regions show substantial overlap with the brain areas identified in the P-FIT theory^25^ and extended Multiple Demand Network^26^. And the most regions are network hubs that are highly connected with other regions and are crucial for information transmission efficiency within the network^5^. The identification of these regions as critical hubs reinforces the notion that differences in intelligence may arise from variations in the brain’s ability to integrate information across distributed regions. Our findings provide further evidence that WM connectivity serves as a key biological substrate for the efficient integration of cognitive processes.

Furthermore, the shared genetic mechanism between WM connectome efficiency and intelligence is not solely attributed to horizontal pleiotropy but may also result from a causal pathway - gene ➔ network efficiency ➔ intelligenc. It was supported by our bidirectional two-sample MR analyses, in which we identified a causal effect from WM connectome efficiency to intelligence, while the reverse causal effect was not significant. This causal pathway is consistent with theories that intelligence arises from the integration of information across distributed brain networks^25,66^.

Using the INT-PGS, the mediation models where Eg of WM connectomes serves as a mediator in the relationship between INT-PGS and intelligence validated the causal pathway - gene ➔ network efficiency ➔ intelligenc. However, the mediation effect accounts for only a small proportion of the total effect. This suggests that the biological pathways through which genes regulate intelligence are highly complex. Our findings align with previous research showing that other brain metrics, such as cortical thickness and surface area, also mediate the effect of genes on intelligence^67^. By identifying network efficiency as one piece of this puzzle, our study contributes to a broader understanding of the neural mechanisms underlying intelligence, reinforcing the idea that intelligence is shaped by multiple interacting factors. Additionally, the finding that combining WM-PGS and NE improved the prediction of intelligence, along with the observed correlation between INT-PGS and Eg of WM connectomes in the mediation model, provides evidence that the genetic factors underlying WM connectome efficiency and intelligence each contain genetic components of the other. This also provides additional evidence from another perspective that WM connectome efficiency and intelligence share a common genetic architecture.

There are several limitations that is necessary to be considered when interpreting the results presented. First, we limited our genomic analyses to participants of white British ancestry. Although this strategy is able to avoid false positive discoveries attributed to population stratifications, it may lead to our results not being generalizable to other populations. Second, the participants included in this study age from 46 to 82, in which efficiency of structural networks decreased with age. Thus, our results may not generalize to other age groups. Third, we did not consider rare genetic variants with population frequencies below 0.01. This may hinder the discovery of additional shared genetic mechanism between topology of structural networks and intelligence. Future analyses based on exome and genome sequence data are necessary.

In summary, our results provided genetic evidence for the association between WM connectome topological properties with intelligence, and revealed the relationship among genes, WM connectome topological properties and intelligence. Our exploration of multiple levels enhanced the understanding of how genes influence intelligence by affecting human brain and highlighted more multi-level studies will be necessary to investigate the neural mechanism of cognition.

## Acknowledgements

This research has been conducted using the UK Biobank Resource under Application Number 49749. This work was supported by the STI2030-Major Projects (2022ZD0213300), National Natural Science Foundation of China (32271145, 81871425, 210510238), Open Research Fund of the State Key Laboratory of Cognitive Neuroscience and Learning (CNLZD2101, CNLZD2303), Fundamental Research Funds for the Central Universities (2017XTCX04).

## Methods

### Participants

The UK Biobank (UKB) is a UK based prospective cohort of ∼500,000 individuals^33^. The study was conducted under UKB application 49749. Ethical approval was obtained from the National Research Ethics Service Committee North West-Haydock (reference 11/NW/0382), and all participants provided written informed consent.

Dataset 1: We utilized initial visit diffusion-weighted magnetic resonance imaging (dMRI) data from ∼40,000 individuals in the UKB, along with T1-weighted MRI data and genotype data from the same participants. After quality control procedures (described below), our final sample included 26,655 participants of unrelated white British ancestry for the primary white matter (WM) genome-wide association study (GWAS) analysis (13,907 female, 12,748 male; age range 46-82 years; mean age 64.2). To replicate the WM GWAS findings, we divided the dataset into two half-sample subsets for replication WM GWAS analysis. The first subset (dataset1-1) included 13,327 participants (6,957 females and 6,370 males; age range 46-82; mean age 64.2). The second subset (dataset1-2) consisted of 13,328 participants (6,950 female and 6,378 male; age range 46-82; mean age 64.3).

Dataset 2: Participants from both the first and second visits who had completed *Fluid intelligence* tests on the touchscreen questionnaire (data field 20016) or the online questionnaires (data field 20191) were drawn from UKB. Participants have up to two minutes to answer as many of the 13 verbal/numerical reasoning questions as possible. For example, one question might be: “Bud is to flower as child is to?” with answer options: “grow/develop/improve/adult/old/do not know/prefer not to answer.” Another question could be: “150…137…125…114…104…What comes next?” with answer options: “96/95/94/93/92/do not know/prefer not to answer.” The fluid intelligence score is calculated as the sum of correct answers given to the 13 questions. After applying quality control procedures (described below) and excluding individuals used in WM GWAS, our final sample of unrelated white British individuals for intelligence GWAS was N = 182,798.

Dataset 3: An independent sample of 4,249 unrelated individuals of non-white-British ancestry was also drawn from UKB for polygenic scores creation and prediction analysis (2,364 female, 1,885 male; age range 45-82; mean age 62.7).

### Genotyping and quality control

We downloaded imputed single nucleotide polymorphism (SNP) data from UKB resource. We first excluded individuals with mismatched self-reported (data field 31) and genetic sex (data field 22,001), those with sex chromosome aneuploidy (data field 22,019), outliers in heterozygosity and missing rates (data field 22,027), and those with ten or more third-degree relatives (data field 22,021). Our primary analyses focused on the White British Unrelated (WBU) subgroup. This subgroup is the intersection of two sample groups defined by Bycroft et al.^33^ the ’White British ancestry’ group (data field 22006) and the ’used in genetic principal components’ group (data field 22020), the latter consisting of high-quality samples filtered to avoid closely related individuals. The genotype quality control was performed using PLINK^68^. In the autosomes, we first removed multi-allele and duplicate variants. Then, we included SNPs with a minor allele frequency (MAF) > 0.01, genotyping rate > 95%, and those passing Hardy–Weinberg equilibrium test (p value > 1 × 10^−6^). Additionally, we excluded variants with an imputation information (INFO) score < 0.8, which is a measurement of imputation quality provided by UKB (resource 1967). The process above resulted in 7,301,482 variants for WM GWAS and 7,269,906 variants for INT GWAS.

The same sample and genotype quality control process was applied to obtain an independent sample of unrelated individuals of non-white-British ancestry, except for the selection criteria in ’White British ancestry’ (data field 22006). The process resulted in 6,970,647 variants.

### Neuroimaging phenotypes

T1-weighted MRI and diffusion-weighted MRI were used for WM structural network construction. The UKB performed pre-processing on MRI data with FMRIB Software Library. The diffusion-weighted MRI preprocessing included eddy current correction, head movement correction, gradient distortion correction, and fiber orientation estimation with bedpostx (Bayesian Estimation of Diffusion Parameters Obtained using Sampling Techniques)^69^. Image acquisition and processing details were summarized in the Supplemental Methods.

To construct the WM connectome, we registered the Brainnetome Atlas^34^ with 246 parcellations defined in standard Montreal Neuroimaging Institute (MNI) space and the WM mask in the T1 space to each participant’s naive diffusion space. Then, we used Camino^70^ (http://camino.cs.ucl.ac.uk/) for deterministic WM fiber tracking with a ball-and-stick model estimated from bedpostx. We utilized nearest-neighbor interpolation and the fourth-order Runge-Kutta method as the tracking algorithm, with a tracking step size set to 2mm. To maintain biological plausibility, we discarded compartments with a mean volume fraction below 0.1 and terminated tracking if curvature exceeded 45 degrees at each 5 mm interval. We further post-processed the tracking results by removing fibers with a length below 20mm or above 250mm. Based on Brainnetome atlas and deterministic tractography results, a fiber number weighted network was constructed for each participant. We excluded 2,052 participants whose images either failed the pipeline or did not pass the quality control, which included visual inspection of registration and tracking.

To describe the topological organization of the WM structural networks, various topological properties at both global and regional level were evaluated. Global characteristics include global efficiency (Eg), local efficiency (Eloc), clustering coefficient (Cp), shortest path length (Lp), and small-world parameters (γ, λ, σ). Regional characteristics include nodal efficiency (NE), nodal local efficiency (NLE), nodal clustering coefficient (NCP), nodal shortest path length (NLP). Definitions of these network topological properties can be found in the Supplemental Methods.

### SNP heritability and genome-wide association analysis

SNP heritability of WM connectome properties was estimated using the genome-based restricted maximum likelihood (GREML)^71^ method, implemented in the Genome-wide complex trait analysis (GCTA) software^72^. Covariates included continuous variables such as age (at imaging), age-squared, and the first 10 genetic principal components, as well as categorical variables including sex, assessment center, and genotype measurement batch. The heritability of nodal properties was considered significant after applying a false discovery rate (FDR) correction across 246 regions.

We performed GWAS for each network property separately using linear regression models with PLINK^68^. The same covariates as in the heritability analysis were regressed out. Statistical significance for SNPs was defined as *P* < 5×10^-8^, while significance for GWAS across 246 regions was defined as *P* < 2×10^-10^ after applying Bonferroni correction. For the GWAS of intelligence, the same covariates were regressed out, excluding the assessment center.

### Identification of independent SNPs and genes

SNPs with genome-wide significance that have LD r^2^ < 0.6 with any other SNPs were identified as independent significant SNPs (Ind. SNPs). Ind. SNPs and all SNPs in LD with them were annotated for gene functional consequences by ANNOVAR^73^. The annotated SNPs were mapped to protein-coding genes based on their physical position on the genome (within 10kb), eQTL associations (using data from the CommonMind Consortium^74^, BRAINEAC^75^, and GTEx v8 brain^76^), and chromatin interaction mapping (using built-in chromatin interaction data for dorsolateral prefrontal cortex and hippocampus^77^, and annotate enhancer/ promoter regions from E053–E082 brain^78^). Default values were used for all other parameters in FUMA. Moreover, genome-wide gene-based association study was performed using MAGMA, implemented in FUMA with default parameters. This process examines the joint association signals of all SNPs within a given gene (including 50 kb upstream to 50 kb downstream of the gene), while considering the LD between the SNPs. Input SNPs were mapped to 19,279 protein-coding genes, and genome-wide significance defined at *P* < 0.05/19279 = 2.593×10^-6^.

### Gene set enrichment analysis

We input a list of genes, which included those mapped from SNPs and significant genes (*P* < 2.593×10^-6^) from the gene-based association analysis, into the Enrichr platform (https://maayanlab.cloud/Enrichr/). The genes were tested against Gene Ontology (GO) gene sets obtained from MsigDB^37^ using a hypergeometric test. The set of background genes consisted of protein-coding genes. FDR multiple testing correction was performed for each data source of tested gene sets.

Additionally, we assessed enrichment in drug targets using the GREP (Genome for Repositioning)^79^ software tool. Using the same list of genes as input, GREP performs Fisher’s exact tests to examine the enrichment of gene sets in drugs of specific Anatomical Therapeutic Chemical (ATC) classes or ICD10 (10th revision of the International Statistical Classification of Diseases and Related Health Problems) disease categories. The target genes for approved or investigated drugs were sourced from two major drug databases, Drug Bank^80^ and Therapeutic Target Database^81^.

### Phenotypic and genetic correlation estimation

We used partial correlation analysis to estimate pairwise phenotypic associations in WBU individuals who had both dMRI and cognitive data (n = 24,704), controlling for sex, age and assessment center. To estimate pairwise genetic correlations between WM connectome properties and intelligence, we employed LD-score regression (LDSC, https://github.com/bulik/ldsc) using summary statistics of univariate GWAS. In LDSC, we utilized the precomputed LD scores from the European population of the 1000 Genomes Project as a reference panel. To uncover the brain-wide correlations, we conducted phenotypic and genetic correlations between regional properties of WM connectome and intelligence.

### Shared genetic variant with colocalization analysis

We explored whether genome-wide significant WM risk loci (independent significant SNPs) were associated with intelligence within a 250 Kb window around each independent SNP. We performed a Bayesian colocalization analysis (COLOC)^38^ to search for evidence for a single causal variant between nodal efficiency and intelligence. COLOC tests five hypotheses: H0, no association with either trait (PP0); H1 or H2, association with either trait 1 or trait 2 (PP1 or PP2); H3, two independent SNPs associated with trait 1 and trait 2 (PP3); H4, one shared SNP associated with both traits (PP4). The analysis generates a posterior probability (PP) for each hypothesis, indicating the degree of support for each. We considered PP4 > 75% as evidence supporting a single causal variant for both traits.

### Mendelian randomization analyses

We evaluated the causal relationships between WM connectome efficiency and intelligence using Mendelian randomization (MR) analysis, considering both global and regional level of the network.

From the exposure GWAS summary statistics, independent significant SNPs were extracted at a significance level of 5×10^-8^. To minimize confounding, we excluded SNPs associated with confounders that have been previously reported to influence both WM connectome^82^ and intelligence^24^. Specifically, we accounted for four potential confounders: education, diabetes, hypertension, and alcohol consumption. Significant associations with these confounders were identified using the NHGRI-EBI GWAS Catalog^39^ (https://www.ebi.ac.uk/gwas/). In addition, SNPs shared by WM connectome efficiency and intelligence were removed based on results from Bayesian colocalization analysis.

Subsequent MR analyses were conducted using TwoSampleMR package^83,84^ (https://mrcieu.github.io/TwoSampleMR/). The harmonization procedure was applied to ensure correct allele alignment, guaranteeing that the selected genetic variants for the exposure and outcome corresponded to the same alleles. We employed nine commonly used MR methods to evaluate the causal associations: median-based methods (simple median, weighted median, and penalized weighted median), inverse variance weighted (IVW) models (fixed effect IVW and multiple random effect IVW), mode-based methods (simple mode, simple mode (NOME), weighed mode, and weighted mode (NOME)).

To evaluate the robustness and reliability of the MR analysis results, we conducted a series of sensitivity analyses on the significant findings. First, leave-one-out analysis was performed to assess whether the causal association is driven by a single SNP. The analysis was implemented using the “*mr_leaveoneout*” function from the TwoSampleMR package. Second, the MR-Egger intercept test was applied to detect horizontal pleiotropy. In MR-Egger regression, the intercept represents the average pleiotropic effect of all genetic variants. A significant non-zero intercept (P < 0.05) indicates the presence of pleiotropy, suggesting that the instrumental variables may affect the outcome through pathways unrelated to the exposure. The test was implemented using the “*mr_pleiotropy_test*” function from the TwoSampleMR package. Third, heterogeneity tests were conducted using the IVW and MR-Egger approaches. Significant heterogeneity (P < 0.05) indicates inconsistent effects of the instrumental variables on the exposure. The analysis was implemented using the “*mr_heterogeneity*” function from the TwoSampleMR package. Additionally, MR estimates using individual SNPs were used as an illustration of heterogeneity, calculated using the Wald ratio method. Significant results were reported after adjusting for FDR correction.

After performing all the MR analyses between nodal efficiency and intelligence, we excluded results estimated with fewer than 6 genetic variants^85^. We prioritized significant results after Bonferroni correction (*P* < 1.1 × 10^-5^, correcting for 246 regions, 2 causal directions, and 9 MR methods). To increase the robustness of our findings, we focused on interpreting MR results that met two criteria: 1) significance in the basic fixed-effect IVW model, and 2) significance in at least four additional MR methods.

### Polygenic scores

Based on GWAS results of intelligence, we constructed polygenic score for intelligence (INT-PGS) in a sample of 26,655 WBU individuals (dataset 1). After quality control, 7,301,482 SNPs were available for PGS calculation (see Genotyping and Quality Control). Based on GWAS results of nodal efficiency, we constructed polygenic scores for each region of WM connectome (WM-PGSs) in an independent sample of 4,249 unrelated individuals of non-white-British ancestry (dataset 3). After quality control, 6,970,647 SNPs were available for PGS calculation. We utilized PRSice-2^86^, a software that automatically performs SNP clumping, PGS calculation, and p-value thresholding. SNPs were clumped and the most significant SNPs with *r^2^* > 0.2 per 250 kb linkage disequilibrium block were retained. PGS for each participant was calculated by summing the clumped SNPs weighted by their effect sizes derived from GWAS summary statistics of White British individuals (GWAS of WM connectome properties and GWAS of cognitive functions). We selected six different p-value thresholds (*P* ≤ 5×10^-8^, 5×10^-4^, 0.005, 0.05, 0.5, 1) for SNP selection in the construction of PGS. Consequently, six PGSs were generated, and we reported the best prediction accuracy achieved by one of these PGSs. The association between PGS and phenotype was estimated in linear models, controlling for the effects of age, age-squared, sex, assessment center (not used in INT-PGS), genotype measurement batch and the first 10 genetic principal components. The additional phenotypic variation explained by PGS (*R^2^*) was used to measure prediction accuracy.

### Mediation analysis

We performed mediation analysis to determine if the effect of INT-PGS on intelligence could be explained by WM connectome efficiency. The effects of age, sex, and assessment center were included as covariates in the direct and indirect paths. The 95% confidence intervals (CIs) were obtained using bootstrapping procedures with 5,000 resamples. An indirect effect was considered significant if the bootstrapped CI did not include zero. In addition to global efficiency of the WM connectome, nodal efficiency was considered as a mediator, one region at a time, to examine the mediation effect of different brain regions. All mediation analysis was carried out using pingouin package^87^.

### Prediction analysis

We performed intelligence prediction using nodal efficiency and its PGS across 246 regions. Partial least squares regression, a widely used algorithm in multivariate neuroimaging research, was adopted for our prediction analysis. We used WM-PGSs, nodal efficiency, and the combination of both as features, respectively. A nested 3-fold cross-validation (CV) approach, incorporating both outer CV and inner CV, was implemented. Specifically, the optimal parameter (the number of components: [2,10] with a step of 1) was selected during the inner CV and utilized in the outer loop for prediction. In the outer CV, the dataset was randomly divided into three folds: two folds were used for training the model, and the remaining fold was used for testing. Pearson’s correlation coefficient between the predicted and actual values was computed to define predictive accuracy. This process was repeated three times, generating three predictive models, and the average predictive accuracy across these three repetitions was taken as the accuracy of the final model. To ensure stable predictive performance, we repeated the above prediction pipeline 100 times, generating 100 Pearson’s correlation coefficients, which were used to compare the predictive accuracy of different feature sets using a t-test.

## Data availability

Our GWAS summary statistics are available upon request. The individual-level data used in the present study were obtained from UKB (https://www.ukbiobank.ac.uk/) under Application Number 49749.

## Code availability

The tractography analysis was conducted using Camino (http://camino.cs.ucl.ac.uk/). Genotype data processing and analysis were carried out with publicly available tools from<colcnt=3>

PLINK (https://www.cog-genomics.org/plink/),

GCTA

(http://cnsgenomics.com/software/gcta/), FUMA

(https://fuma.ctglab.nl/),

MAGMA

(https://ctg. cncr.nl/software/magma), Enrichr (https://maayanlab.cloud/Enrichr/), GREP (https://github.com/saorisakaue/GREP), LD-Score Regression (https://github.com/ bulik/ldsc/), PRSice-2 (https://choishingwan.github.io/PRSice/), coloc (https://chr1swallace.github.io/coloc/) R package, TwoSampleMR (https://mrcieu.github.io/

TwoSampleMR/) R package, pingouin (https://pingouin-stats.org/) Python package, and scikit-learn (https://scikit-learn.org/) Python package. Custom codes for the analyses in this paper are available through GitHub (https://github.com/Sheeya-Dong/WM_intelligence).

## References

1. Mansour L, S., Tian, Y., Yeo, B. T. T., Cropley, V. & Zalesky, A. High-resolution connectomic fingerprints: Mapping neural identity and behavior. NeuroImage 229, 117695 (2021).

2. Bullmore, E. & Sporns, O. Complex brain networks: graph theoretical analysis of structural and functional systems. Nat. Rev. Neurosci. 10, 186–198 (2009).

3. Liao, X., Vasilakos, A. V. & He, Y. Small-world human brain networks: Perspectives and challenges. Neurosci. Biobehav. Rev. 77, 286–300 (2017).

4. Park, H.-J. & Friston, K. Structural and Functional Brain Networks: From Connections to Cognition. Science 342, 1238411 (2013).

5. Heuvel, M. P. van den & Sporns, O. Rich-Club Organization of the Human Connectome. J. Neurosci. 31, 15775–15786 (2011).

6. Shu, N., Wang, X., Bi, Q., Zhao, T. & Han, Y. Disrupted Topologic Efficiency of White Matter Structural Connectome in Individuals with Subjective Cognitive Decline. Radiology 286, 229–238 (2018).

7. Chen, Q. et al. Increased segregation of structural brain networks underpins enhanced broad cognitive abilities of cognitive training. Hum. Brain Mapp. 42, 3202–3215 (2021).

8. Iturria-Medina, Y., Sotero, R. C., Canales-Rodríguez, E. J., Alemán-Gómez, Y. & Melie-García, L. Studying the human brain anatomical network via diffusion-weighted MRI and Graph Theory. NeuroImage 40, 1064–1076 (2008).

9. Córdova-Palomera, A. et al. Genetic control of variability in subcortical and intracranial volumes. Mol. Psychiatry 26, 3876–3883 (2021).

10. van der Meer, D. et al. The genetic architecture of human cortical folding. Sci. Adv. 7, eabj9446 (2021).

11. Grasby, K. L. et al. The genetic architecture of the human cerebral cortex. Science 367, eaay6690 (2020).

12. Zhao, B. et al. Common variants contribute to intrinsic human brain functional networks. Nat. Genet. 54, 508–517 (2022).

13. Zhao, B. et al. Common genetic variation influencing human white matter microstructure. Science 372, eabf3736 (2021).

14. Sha, Z., Schijven, D., Fisher, S. E. & Francks, C. Genetic architecture of the white matter connectome of the human brain. Sci. Adv. 9, eadd2870 (2023).

15. Wainberg, M. et al. Genetic architecture of the structural connectome. Nat. Commun. 15, 1962 (2024).

16. Bullmore, E. & Sporns, O. The economy of brain network organization. Nat. Rev. Neurosci. 13, 336–349 (2012).

17. Zhao, T. et al. Test-retest reliability of white matter structural brain networks: a multiband diffusion MRI study. Front. Hum. Neurosci. 9, 59 (2015).

18. Deary, I. J. The stability of intelligence from childhood to old age. Curr. Dir. Psychol. Sci. 23, 239–245 (2014).

19. Deary, I. J., Strand, S., Smith, P. & Fernandes, C. Intelligence and educational achievement. Intelligence 35, 13–21 (2007).

20. Strenze, T. Intelligence and socioeconomic success: A meta-analytic review of longitudinal research. Intelligence 35, 401–426 (2007).

21. Deary, I. J. et al. Intergenerational social mobility and mid-life status attainment: Influences of childhood intelligence, childhood social factors, and education. Intelligence 33, 455–472 (2005).

22. Wraw, C., Deary, I. J., Gale, C. R. & Der, G. Intelligence in youth and health at age 50. Intelligence 53, 23–32 (2015).

23. Cm, C. et al. Childhood intelligence in relation to major causes of death in 68 year follow-up: prospective population study. BMJ 357, (2017).

24. Deary, I. J., Cox, S. R. & Hill, W. D. Genetic variation, brain, and intelligence differences. Mol. Psychiatry 27, 335–353 (2022).

25. Jung, R. E. & Haier, R. J. The Parieto-Frontal Integration Theory (P-FIT) of intelligence: converging neuroimaging evidence. Behav. Brain Sci. 30, 135–154; discussion 154-187 (2007).

26. Camilleri, J. A. et al. Definition and characterization of an extended multiple-demand network. NeuroImage 165, 138–147 (2018).

27. S, K., et al. Positive association between cognitive ability and cortical thickness in a representative US sample of healthy 6 to 18 year-olds. Intelligence 37, 145–155 (2009).

28. Dubois, J., Galdi, P., Paul, L. K. & Adolphs, R. A distributed brain network predicts general intelligence from resting-state human neuroimaging data. Philos. Trans. R. Soc. Lond. B. Biol. Sci. 373, 20170284 (2018).

29. Basten, U., Hilger, K. & Fiebach, C. J. Where smart brains are different: A quantitative meta-analysis of functional and structural brain imaging studies on intelligence. Intelligence 51, 10–27 (2015).

30. Hill, W. D. et al. A combined analysis of genetically correlated traits identifies 187 loci and a role for neurogenesis and myelination in intelligence. Mol. Psychiatry 24, 169–181 (2019).

31. G, D., et al. Study of 300,486 individuals identifies 148 independent genetic loci influencing general cognitive function. Nat. Commun. 9, (2018).

32. Savage, J. E. et al. Genome-wide association meta-analysis in 269,867 individuals identifies new genetic and functional links to intelligence. Nat. Genet. 50, 912–919 (2018).

33. Bycroft, C. et al. The UK Biobank resource with deep phenotyping and genomic data. Nature 562, 203–209 (2018).

34. Fan, L. et al. The Human Brainnetome Atlas: A New Brain Atlas Based on Connectional Architecture. Cereb. Cortex 26, 3508–3526 (2016).

35. Leeuw, C. A. de, Mooij, J. M., Heskes, T. & Posthuma, D. MAGMA: Generalized Gene-Set Analysis of GWAS Data. PLOS Comput. Biol. 11, e1004219 (2015).

36. Watanabe, K., Taskesen, E., van Bochoven, A. & Posthuma, D. Functional mapping and annotation of genetic associations with FUMA. Nat. Commun. 8, 1826 (2017).

37. Liberzon, A. et al. Molecular signatures database (MSigDB) 3.0. Bioinformatics 27, 1739–1740 (2011).

38. Giambartolomei, C. et al. Bayesian test for colocalisation between pairs of genetic association studies using summary statistics. PLoS Genet. 10, e1004383 (2014).

39. Buniello, A. et al. The NHGRI-EBI GWAS Catalog of published genome-wide association studies, targeted arrays and summary statistics 2019. Nucleic Acids Res. 47, D1005–D1012 (2019).

40. Tzourio-Mazoyer, N. et al. Automated anatomical labeling of activations in SPM using a macroscopic anatomical parcellation of the MNI MRI single-subject brain. NeuroImage 15, 273–289 (2002).

41. Arnatkeviciute, A. et al. Genetic influences on hub connectivity of the human connectome. Nat. Commun. 12, 4237 (2021).

42. Bc, C., R, W. & Eb, L. Neurodegenerative Disease Tauopathies. Annu. Rev. Pathol. 19, (2024).

43. Zhao, B. et al. Large-scale GWAS reveals genetic architecture of brain white matter microstructure and genetic overlap with cognitive and mental health traits (n[=[17,706). Mol. Psychiatry 26, 3943–3955 (2021).

44. C, M., et al. Larger cerebral cortex is genetically correlated with greater frontal area and dorsal thickness. Proc. Natl. Acad. Sci. U. S. A. 120, (2023).

45. Zhao, B. et al. Genome-wide association analysis of 19,629 individuals identifies variants influencing regional brain volumes and refines their genetic co-architecture with cognitive and mental health traits. Nat. Genet. 51, 1637–1644 (2019).

46. Noel, J. et al. Surface Expression of AMPA Receptors in Hippocampal Neurons Is Regulated by an NSF-Dependent Mechanism. Neuron 23, 365–376 (1999).

47. Srivastava, T. et al. A TLR/AKT/FoxO3 immune tolerance–like pathway disrupts the repair capacity of oligodendrocyte progenitors. J. Clin. Invest. 128, 2025–2041 (2018).

48. May-Wilson, S. et al. Large-scale GWAS of food liking reveals genetic determinants and genetic correlations with distinct neurophysiological traits. Nat. Commun. 13, 2743 (2022).

49. Elvsåshagen, T. et al. The genetic architecture of the human thalamus and its overlap with ten common brain disorders. Nat. Commun. 12, 2909 (2021).

50. van der Meer, D. et al. Understanding the genetic determinants of the brain with MOSTest. Nat. Commun. 11, 3512 (2020).

51. Smith, S. M. et al. An expanded set of genome-wide association studies of brain imaging phenotypes in UK Biobank. Nat. Neurosci. 24, 737–745 (2021).

52. van der Meer, D. et al. Boosting Schizophrenia Genetics by Utilizing Genetic Overlap With Brain Morphology. Biol. Psychiatry 92, 291–298 (2022).

53. Koenis, M. M. G. et al. Association between structural brain network efficiency and intelligence increases during adolescence. Hum. Brain Mapp. 39, 822–836 (2018).

54. Li, Y. et al. Brain Anatomical Network and Intelligence. PLOS Comput. Biol. 5, e1000395 (2009).

55. Fischer, F. U., Wolf, D., Scheurich, A. & Fellgiebel, A. Association of Structural Global Brain Network Properties with Intelligence in Normal Aging. PLOS ONE 9, e86258 (2014).

56. Zheng, N., Fraenkel, E., Pabo, C. O. & Pavletich, N. P. Structural basis of DNA recognition by the heterodimeric cell cycle transcription factor E2F–DP. Genes Dev. 13, 666–674 (1999).

57. Zhang, Y. & Chellappan, S. P. Cloning and characterization of human DP2, a novel dimerization partner of E2F. Oncogene 10, 2085–2093 (1995).

58. Park, K.-K. et al. Inhibitory effects of novel E2F decoy oligodeoxynucleotides on mesangial cell proliferation by coexpression of E2F/DP. Biochem. Biophys. Res. Commun. 308, 689–697 (2003).

59. Sang, T., Cao, Q., Wang, Y., Liu, F. & Chen, S. Overexpression or silencing of FOXO3a affects proliferation of endothelial progenitor cells and expression of cell cycle regulatory proteins. PloS One 9, e101703 (2014).

60. Marzi, L. et al. FOXO3a and the MAPK p38 are activated by cetuximab to induce cell death and inhibit cell proliferation and their expression predicts cetuximab efficacy in colorectal cancer. Br. J. Cancer 115, 1223–1233 (2016).

61. Liang, C. et al. Serotonin promotes the proliferation of serum-deprived hepatocellular carcinoma cells via upregulation of FOXO3a. Mol. Cancer 12, 14 (2013).

62. Cui, Y. et al. Decreased RNA[binding protein IGF2BP2 downregulates NT5DC2, which suppresses cell proliferation, and induces cell cycle arrest and apoptosis in diffuse large B[cell lymphoma cells by regulating the p53 signaling pathway. Mol. Med. Rep. 26, 286 (2022).

63. Lee, S., Koh, W., Kim, H.-T., Kim, C.-H. & Lee, S. Cancer-upregulated gene 2 (CUG2) overexpression induces apoptosis in SKOV-3 cells. Cell Biochem. Funct. 28, 461–468 (2010).

64. Nishino, T. et al. CENP-T-W-S-X forms a unique centromeric chromatin structure with a histone-like fold. Cell 148, 487–501 (2012).

65. Giannuzzi, G. et al. Evolutionary dynamism of the primate LRRC37 gene family. Genome Res. 23, 46–59 (2013).

66. Duncan, J., Assem, M. & Shashidhara, S. Integrated Intelligence from Distributed Brain Activity. Trends Cogn. Sci. 24, 838–852 (2020).

67. Williams, C. M., Peyre, H. & Ramus, F. Brain volumes, thicknesses, and surface areas as mediators of genetic factors and childhood adversity on intelligence. Cereb. Cortex 33, 5885–5895 (2023).

68. Purcell, S. et al. PLINK: A Tool Set for Whole-Genome Association and Population-Based Linkage Analyses. Am. J. Hum. Genet. 81, 559–575 (2007).

69. Jbabdi, S., Sotiropoulos, S. N., Savio, A. M., Graña, M. & Behrens, T. E. J. Model-based analysis of multishell diffusion MR data for tractography: how to get over fitting problems. Magn. Reson. Med. 68, 1846–1855 (2012).

70. Cook, P. A. et al. Camino: Open-Source Diffusion-MRI Reconstruction and Processing. in (14th scientific meeting of the international society for magnetic resonance in medicine, Seattle WA, USA, 2006).

71. Yang, J. et al. Common SNPs explain a large proportion of the heritability for human height. Nat. Genet. 42, 565–569 (2010).

72. Yang, J., Lee, S. H., Goddard, M. E. & Visscher, P. M. GCTA: A Tool for Genome-wide Complex Trait Analysis. Am. J. Hum. Genet. 88, 76–82 (2011).

73. Wang, K., Li, M. & Hakonarson, H. ANNOVAR: functional annotation of genetic variants from high-throughput sequencing data. Nucleic Acids Res. 38, e164 (2010).

74. Fromer, M. et al. Gene expression elucidates functional impact of polygenic risk for schizophrenia. Nat. Neurosci. 19, 1442–1453 (2016).

75. Ramasamy, A. et al. Genetic variability in the regulation of gene expression in ten regions of the human brain. Nat. Neurosci. 17, 1418–1428 (2014).

76. GTEx Consortium. Human genomics. The Genotype-Tissue Expression (GTEx) pilot analysis: multitissue gene regulation in humans. Science 348, 648–660 (2015).

77. Schmitt, A. D. et al. A Compendium of Chromatin Contact Maps Reveals Spatially Active Regions in the Human Genome. Cell Rep. 17, 2042–2059 (2016).

78. Kundaje, A. et al. Integrative analysis of 111 reference human epigenomes. Nature 518, 317–330 (2015).

79. Sakaue, S. & Okada, Y. GREP: genome for REPositioning drugs. Bioinformatics 35, 3821–3823 (2019).

80. Wishart, D. S. et al. DrugBank 5.0: a major update to the DrugBank database for 2018. Nucleic Acids Res. 46, D1074–D1082 (2018).

81. Li, Y. H. et al. Therapeutic target database update 2018: enriched resource for facilitating bench-to-clinic research of targeted therapeutics. Nucleic Acids Res. 46, D1121–D1127 (2018).

82. Chen, H.-J. et al. Effects of Vascular Risk Factors on the White Matter Network Architecture of the Brain. Neurosci. Bull. (2024) doi:10.1007/s12264-024-01274-3.

83. Hemani, G., Tilling, K. & Smith, G. D. Orienting the causal relationship between imprecisely measured traits using GWAS summary data. PLOS Genet. 13, e1007081 (2017).

84. Hemani, G. et al. The MR-Base platform supports systematic causal inference across the human phenome. eLife 7, e34408 (2018).

85. Zhao, B. et al. Heart-brain connections: Phenotypic and genetic insights from magnetic resonance images. Science 380, abn6598 (2023).

86. Choi, S. W. & O’Reilly, P. F. PRSice-2: Polygenic Risk Score software for biobank-scale data. GigaScience 8, giz082 (2019).

87. Vallat, R. Pingouin: statistics in Python. J. Open Source Softw. 3, 1026 (2018).

