## Supplementary Fig for "Shared genomic architecture of the brain white matter structural connectome and intelligence"

**Supplementary Information**

### Supplementary Methods

#### Image acquisition and processing

Two modalities of MRI scans were used, including T1-weighted MRI and diffusion-weighted MRI. The neuroimaging acquisition and processing pipeline have been thoroughly described elsewhere [1]. T1-weighted MRI was acquired with a 3D magnetization-prepared rapid gradient echo sequence with sagittal scan orientation. The diffusion-weighted MRI scan utilized an echo-planar imaging sequence, a voxel size of 2.0×2.0×2.0 mm, and partial Fourier sampling factors of 6/8. The diffusion sensitizing gradients included 10 b = 0 s/mm² volumes, 50 b = 1 000 s/mm² volumes, and 50 b = 2000 s/mm² volumes, with an echo time of 92 ms and a repetition time of 3600 ms.

UK Biobank performed pre-processing on MRI data with FMRIB Software Library. T1-MRI underwent gradient distortion correction, followed by cropping to optimize the field of view. Subsequently, brain scalping was performed to remove cranial matter, and FMRIB's Automated Segmentation Tool was used to obtain the bias field and components of gray matter, white matter, and cerebrospinal fluid. The volume of each component was then determined.

For diffusion-weighted MRI, preprocessing involved eddy current correction, head movement correction, and gradient distortion correction of the original image in combination with a field map estimated from posterior-to-anterior and anterior-toposterior directed non-diffusional scans. The preprocessed image was then analyzed using the bedpostx tool, which employs model-based spherical deconvolution to estimate up to 3 fiber orientations per voxel. This approach allowed for within-voxel modeling of multi-fiber tract orientation structure, using Bayesian Estimation of Diffusion Parameters Obtained using Sampling Techniques [2].

#### WM network construction and analysis

To construct the WM connectome, we registered the fine Brainnetome Atlas [3] with 246 parcellations defined in standard Montreal Neuroimaging Institute space and the WM mask in the T1 space to each participant’s naive diffusion space. We used Camino [4] (http://camino.cs.ucl.ac.uk/) to perform deterministic WM fiber tracking on the ballstick model in the diffusion space, with the transformed WM mask used as seeding regions. We utilized nearest-neighbor interpolation and the Fourth-order Runge-Kutta method as tracking algorithm, with the tracking step size set to 2mm. To maintain biological plausibility, we discarded compartments with a mean volume fraction below 0.1 and terminated tracking if curvature exceeded 45 degrees at each 5 mm interval. We further post-processed the tracking results by removing fibers with a length below 20mm or above 250mm in the individual diffusion space. Finally, we used the conmat command to construct a fiber number-weighted connectivity network based on the Brainnetome atlas. To ensure accuracy, we had seven experienced colleagues visually cross-check the transformations of the Brainnetome atlas and the plausibility of tracking.

#### Calculation of network topological properties

To describe the topologic organization of the WM structural networks, various topological properties of a network from both global and nodal characteristics were calculated using Gretna [5] (http://www.nitrc.org/projects/gretna/). Global characteristics include global efficiency (Eg), local efficiency (Eloc), clustering coefficient (Cp), shortest path length (Lp), and small-world parameters (γ,λ,σ). Regional characteristics include nodal efficiency (NE), nodal local efficiency (NLE), nodal clustering coefficient (NCP), nodal shortest path length (NLP). The definitions of these properties are as follows.

*Clustering coefficient*

The clustering coefficient $C^{w}$ is calculated as follows:

$$C^{w}(G)= \frac{1}{n} \sum_{i\in G} \frac{2t_{i}^{w}}{k_{i}(k_{i}-1)}= \frac{1}{n} \sum_{i\in G} \frac{\sum_{j,h\in N} {w_{ij}w_{ih}w_{jh}}^{1/3}}{k_{i}(k_{i}-1)}$$

Where $w_{ij}$ is the connection weights between nodes $i$ and $j$, $t_{i}^{w}$ is weighted geometric mean of triangles around $i$, $w_{ij}$ is the connection weight between $i$ and $j$.

The clustering coefficient, Cp, of a network is the average of the clustering coefficient over all nodes and indicates the extent of the local interconnectivity or cliquishness in a network [6].

*Shortest path length*

The shortest path length between $i$ and $j$ is calculated as follows:

$$L_{ij}^{w}= \sum_{a_{uv}\in g_{i\leftrightarrow j}}^{w} f(w_{uv})$$

where $f$ is a map (e.g., an inverse) from weight to length and $g_{i\leftrightarrow j}$ is the shortest weighted path between $i$ and $j$.

The shortest path length of a network was computed as follows:

$$Lp(G)= \frac{1}{n(n-1)}\sum_{i\neq j\in G} L_{ij}^{w}$$

where n is the number of nodes in the network. The Lp of a network quantifies the ability for information to propagate in parallel.

*Small-world parameters*

To examine the small-world properties, the clustering coefficient, Cp, and the shortest path length, Lp, of the brain networks were compared with those of random networks. In this study, we generated 100 matched random networks, which had the same number of nodes, edges, and degree distribution as the real networks [7]. Of note, we retained the weight of each edge during the randomization procedure such that the weight distribution of the network was preserved. Furthermore, we computed the normalized Lp, $\lambda=\frac{{Lp}^{real}}{{Lp}^{rand}}$ , and the normalized Cp, $\gamma=\frac{{Cp}^{real}}{{Cp}^{rand}}$ , where ${Lp}^{rand}$ and ${Cp}^{rand}$ are the mean Lp and the mean Cp of 100 matched random networks, respectively. Importantly, two parameters correct the differences in the edge number and degree distribution of the networks across individuals. A real network would be considered small-world if $\gamma>1$ and $\lambda\approx1$ [6]. Thus, a small-world network not only has a higher local interconnectivity, but it also has an approximately equivalent shortest path length compared with random networks. These two measurements can be summarized into a simple quantitative metric, small-worldness, $\sigma=\frac{\gamma}{\lambda}$ , which is typically greater than 1 for small-world networks [8].

*Efficiency*

The global efficiency of a network measures the global efficiency of the parallel information transfer in the network, which is defined as:

$$E_{glob}\left( G \right)=\frac{1}{n\left( n-1 \right)}\sum_{i\neq j\in G} \frac{1}{L_{ij}}$$

where $L_{ij}$ is the shortest weighted path length between node $i$ and $j$ in $G$.

The local efficiency of $G$ shows how efficient the communication is among the first neighbors of the node $i$ when it is removed, is defined as:

$$E_{loc}\left( G \right)=\frac{1}{n}\sum_{i\in G} E_{glob}\left( G_{i} \right)$$

where $G_{i}$is the subgraph composed of the nearest neighbors of node $i$.

The nodal efficiency of one node measures the average shortest path length between a given node $i$ and all of the other nodes in the network, which is defined as:

$${NE}_{glob}\left( i \right) =\frac{1}{n-1}\sum_{i\neq j\in G} \frac{1}{L_{ij}}$$

where $L_{ij}$ is the shortest weighted path length between node $i$ and $j$ in $G$.

The nodal local efficiency is defined as:

$${NE}_{loc}\left( i \right)=E_{glob}\left( G_{i} \right)$$

where $G_{i}$is the subgraph composed of the nearest neighbors of node $i$.

### List of Supplementary Tables

Table S1. SNP heritability of regional properties of WM structural network

Table S2. Independent significant SNPs associated with global efficiency of WM network

Table S3. Independent significant SNPs associated with nodal efficiency across 246 regions with min-P approach

Table S4. Genes identified by gene-based association analysis for the minimum p-value GWAS of nodal efficiency

Table S5. Genes identified by gene-based association analysis for the minimum p-value GWAS of nodal local efficiency

Table S6. Genes identified by gene-based association analysis for the minimum p-value GWAS of nodal shortest path length

Table S7. Genes identified by gene-based association analysis for the minimum p-value GWAS of nodal clustering coefficient

Table S8. Enriched GO gene sets for the nodal efficiency genes

Table S9. Enriched GO gene sets for the nodal local efficiency genes

Table S10. Enriched GO gene sets for the nodal shortest path length genes

Table S11. Enriched GO gene sets for the nodal clustering coefficient genes

Table S12-1. Phenotypic associations between intelligence and global properties of WM network

Table S12-2. Genetic associations between intelligence and global properties of WM network

Table S13. Independent significant SNPs associated with intelligence

Table S14. Shared genetic variants between global efficiency of WM network and intelligence

Table S15. Shared genetic variants between nodal efficiency of WM network and intelligence

Table S16. Causal relationships between global efficiency and intelligence

Table S17. The significant causal relationships from NE to intelligence

Table S18. Leave-one-out analyses for the significant results of forward MR analyses

Table S19. Horizontal pleiotropy tests for the significant results of forward MR analyses

Table S20. Heterogeneity tests for the significant results of forward MR analyses

Table S21. Causal effects of global efficiency on intelligence using individual SNPs

Table S22. Validating independent significant SNPs of global efficiency in the first half-sample dataset

Table S23. Validating independent significant SNPs of global efficiency in the second half-sample dataset

Table S24. Validating independent significant SNPs of global efficiency in AAL90 atlas

### Supplementary Figures

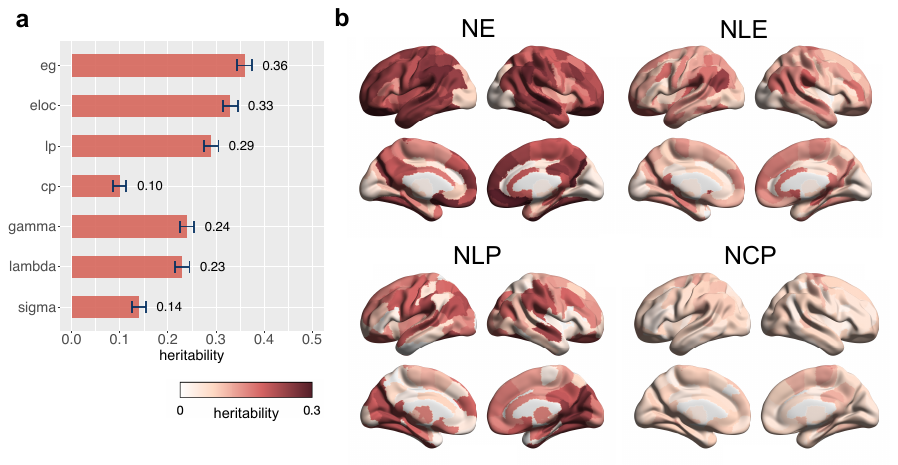

**Supplementary Figure 1.** SNP heritability of global and regional properties of WM structural network. (a) SNP heritability of global properties. The bars and error bars represent the mean and standard deviations of heritability values, respectively. (b) SNP heritability of regional properties across brain regions. WM, white matter; Eg. global efficiency; Eloc, local efficiency; Cp, clustering coefficient; Lp, shortest path length; gamma, lambda, sigma, small-world parameters; NE, nodal efficiency; NLE, nodal local efficiency; NLP, nodal shortest path length; NCP, nodal clustering coefficient.

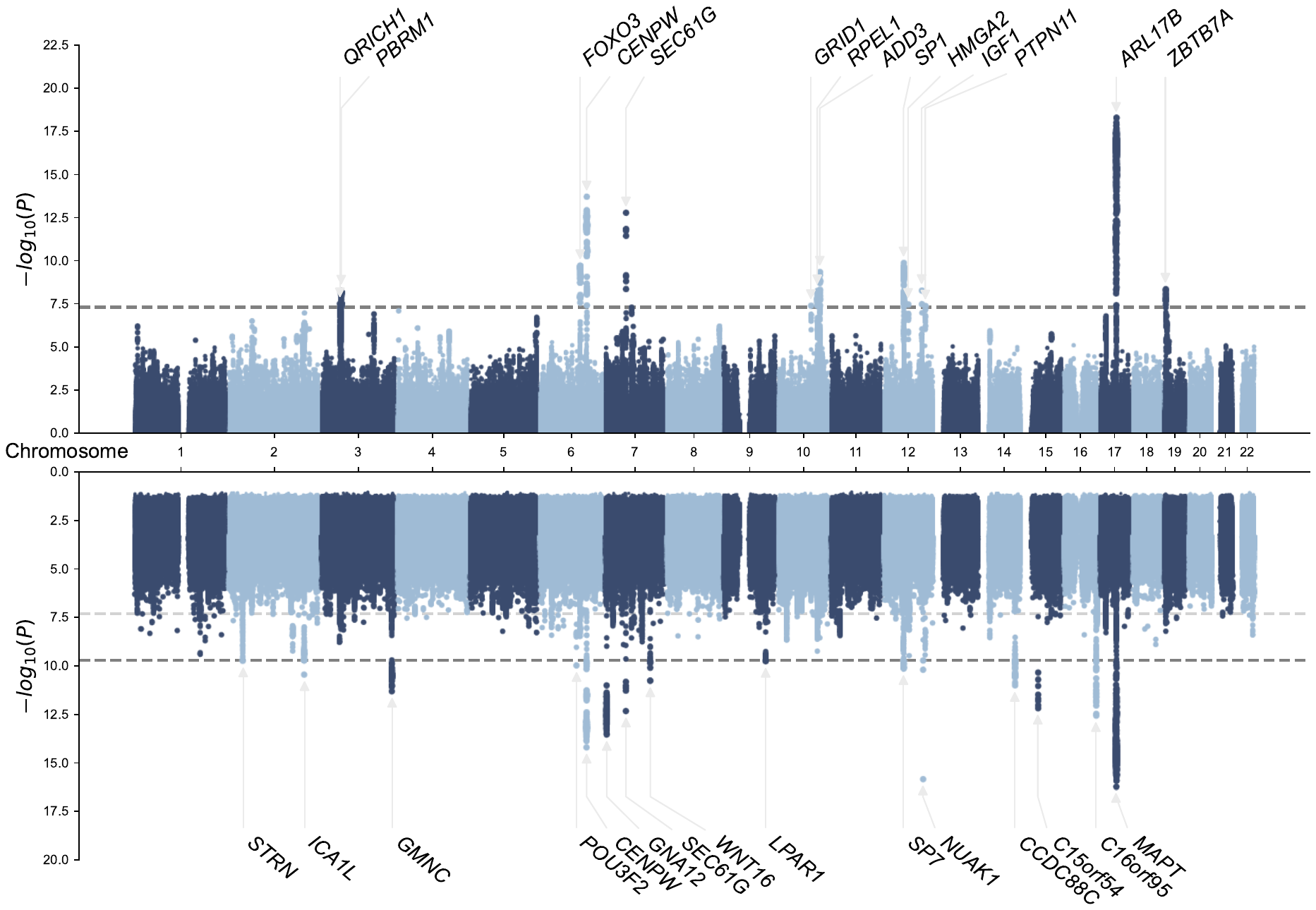

**Supplementary Figure 2.** Manhattan plots with genetic variants identified through univariate GWAS of local efficiency (upper) and multiple univariate GWAS of nodal local efficiency across 246 regions with min-P approach (lower). The grey lines indicate genome-wide significance threshold (P < 5 × 10–8), and Bonferroni-corrected threshold for the multiple univariate GWAS (P < 8.3 × 10–9). The independent significant variants are annotated by their nearest genes.

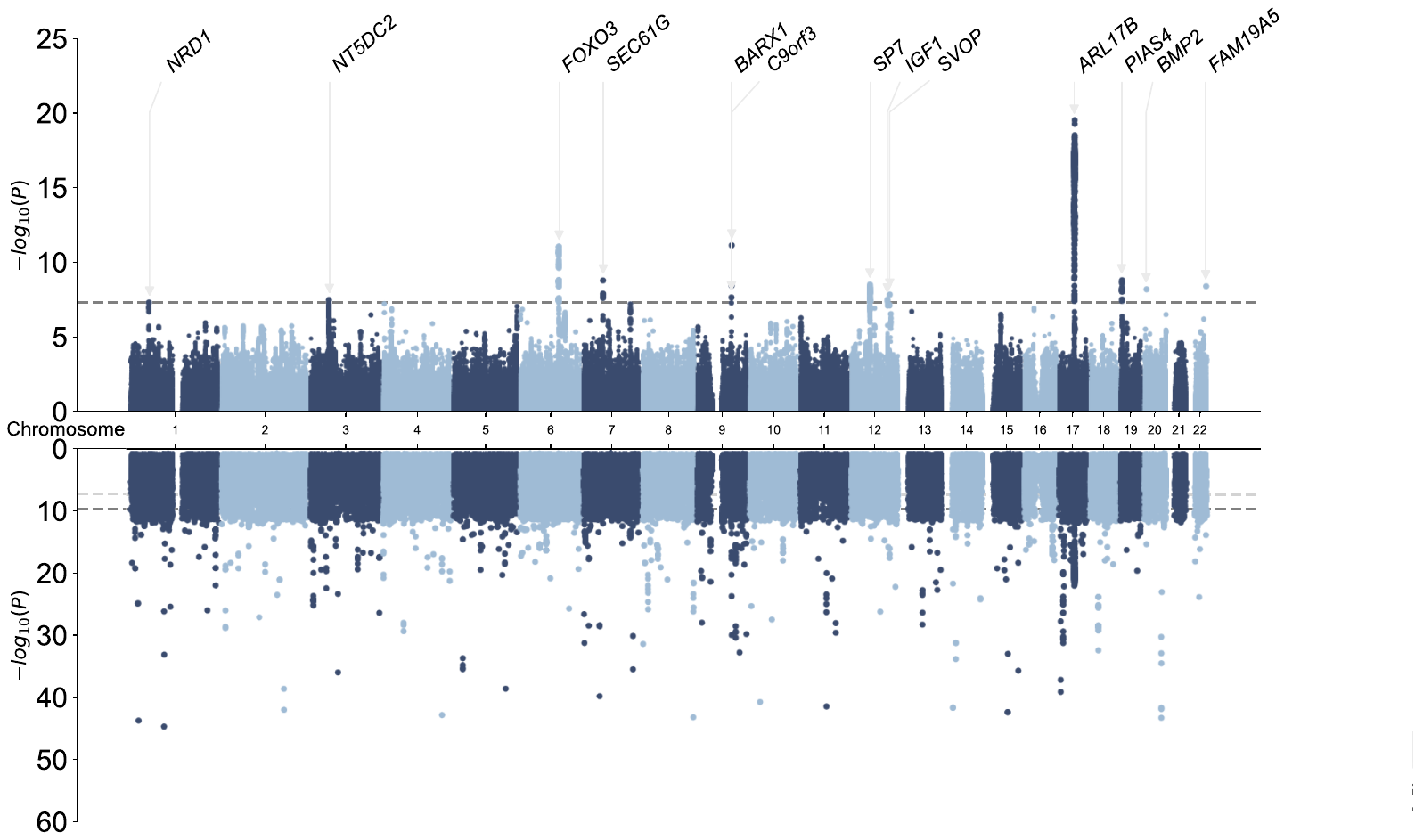

**Supplementary Figure 3.** Manhattan plots with genetic variants identified through univariate GWAS of shortest path length (upper) and multiple univariate GWAS of nodal shortest path length across 246 regions with min-P approach (lower). The grey lines indicate genome-wide significance threshold (P < 5 × 10–8), and Bonferroni-corrected threshold for the multiple univariate GWAS (P < 8.3 × 10–9). The independent significant variants are annotated by their nearest genes (upper).

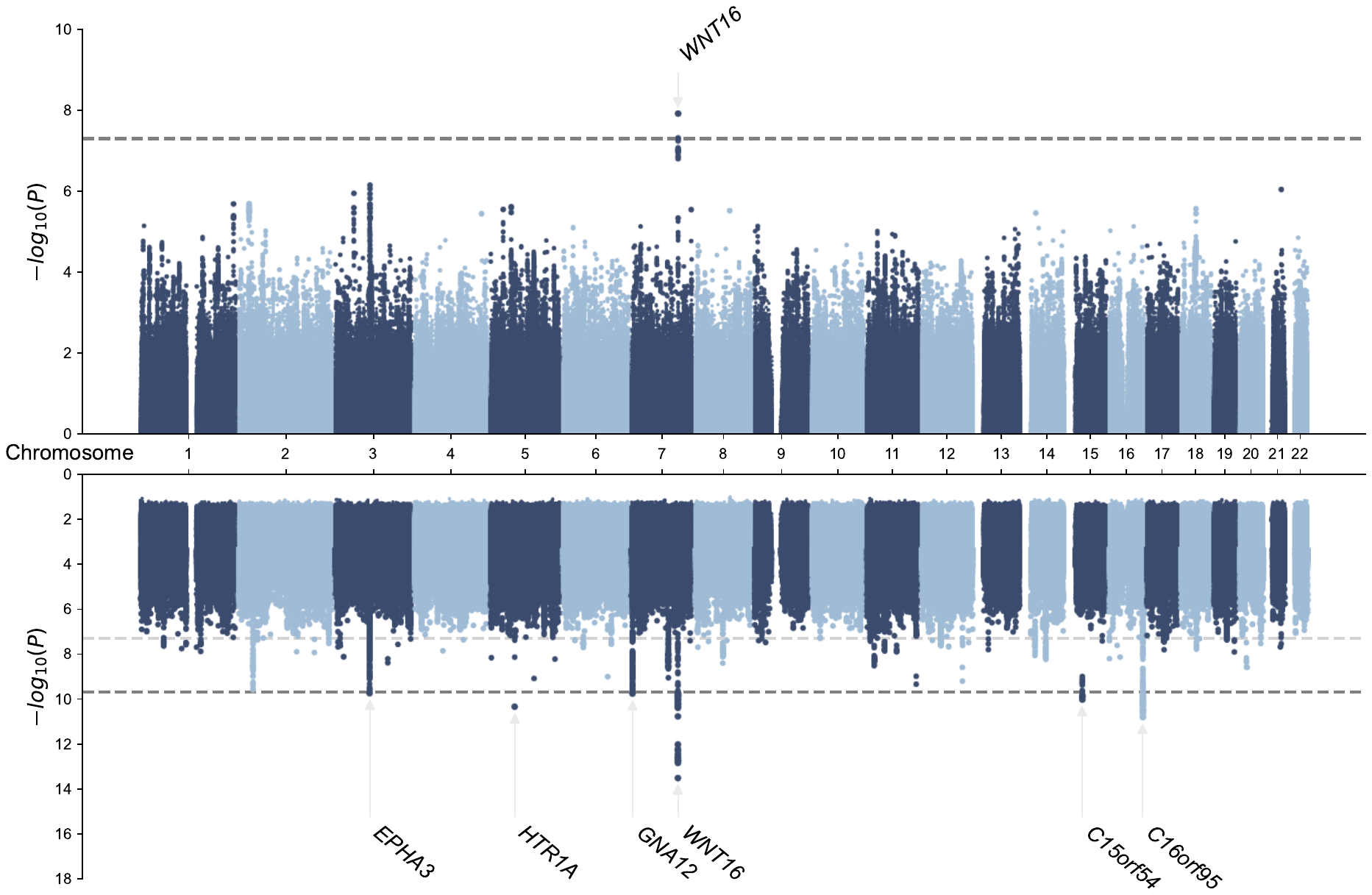

**Supplementary Figure 4.** Manhattan plots with genetic variants identified through univariate GWAS of clustering coefficient (upper) and multiple univariate GWAS of nodal clustering coefficient across 246 regions with min-P approach (lower). The grey lines indicate genome-wide significance threshold (P < 5 × 10–8), and Bonferroni-corrected threshold for the multiple univariate GWAS (P < 8.3 × 10–9). The independent significant variants are annotated by their nearest genes.

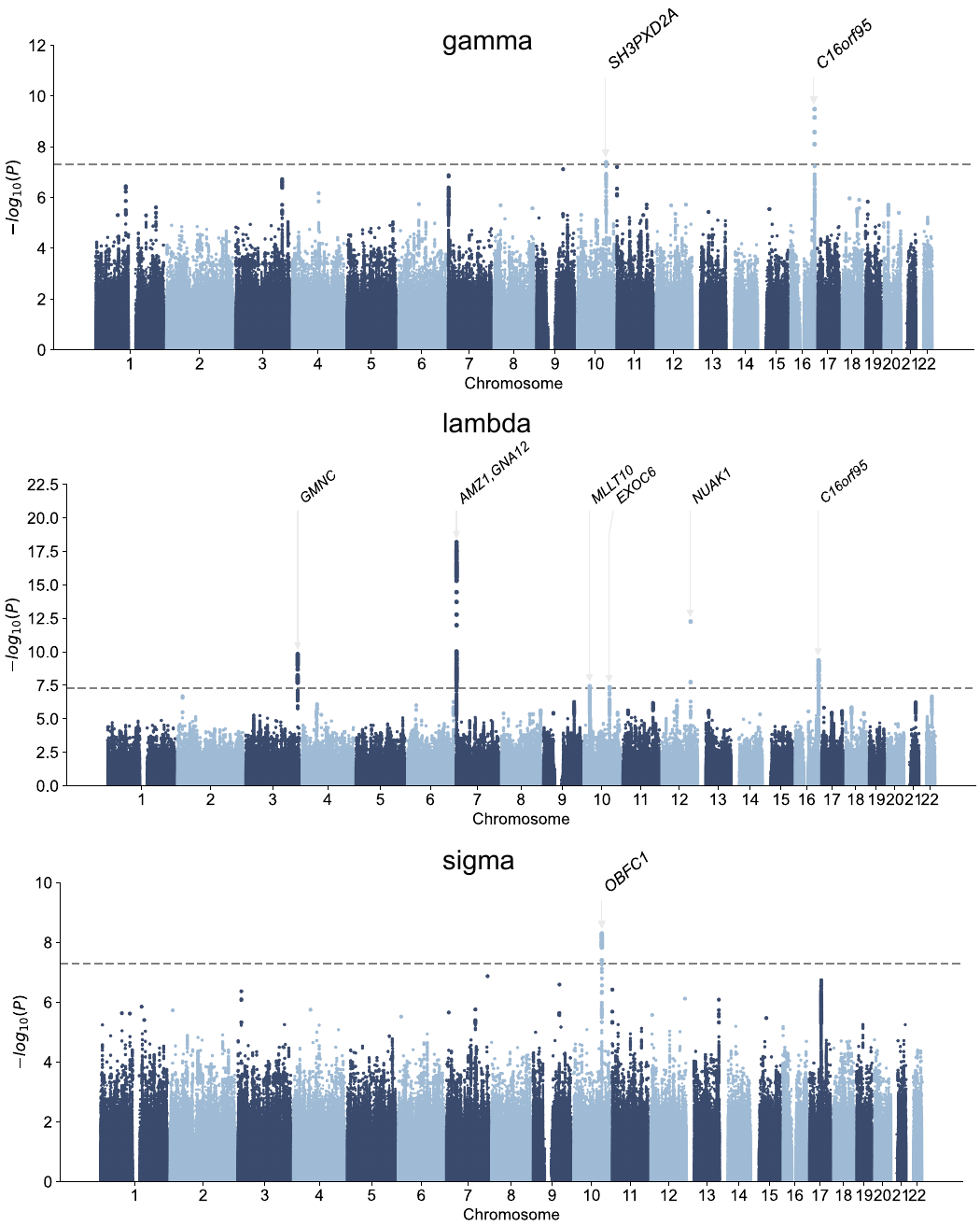

**Supplementary Figure 5.** Manhattan plots with genetic variants identified through univariate GWAS of small-world parameters, gamma, lambda, and sigma. The grey lines indicate genome-wide significance threshold (P < 5 × 10–8). The independent significant variants are annotated by their nearest genes.

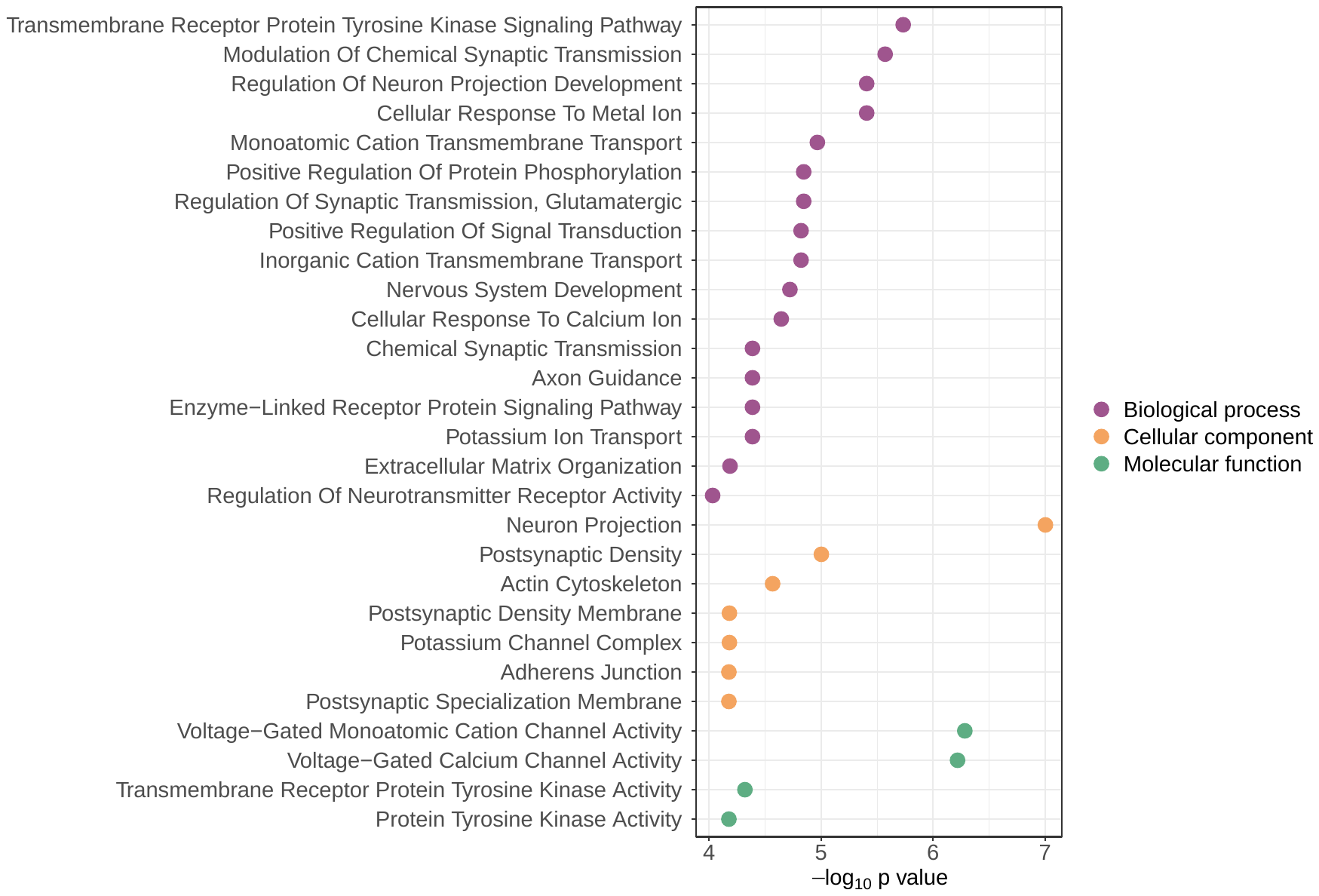

**Supplementary Figure 6.** Top GO gene sets (P < 1 × 10–4) with significant enrichment of association with nodal efficiency across 246 regions.

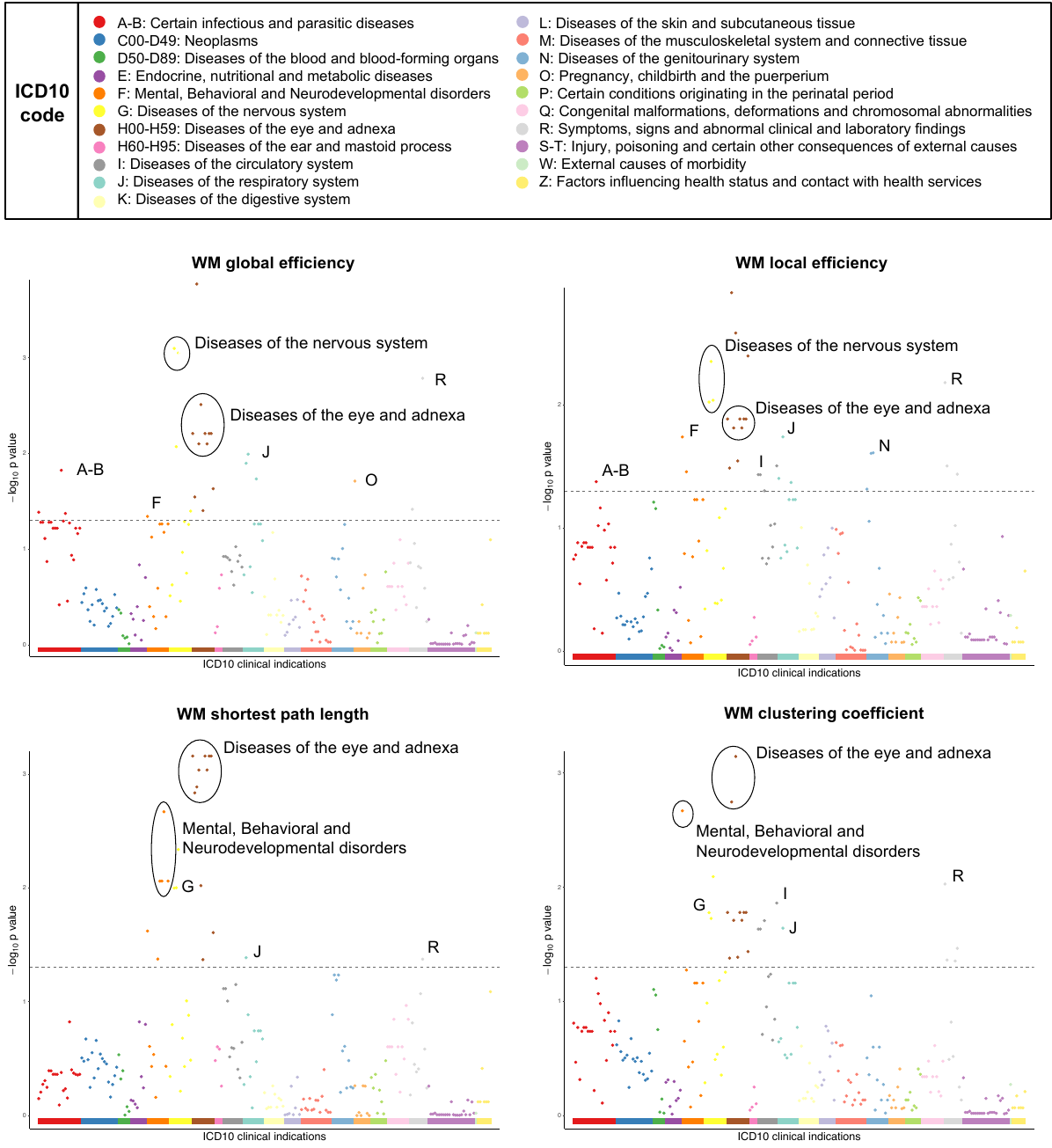

**Supplementary Figure 7.** Enrichment of WM network genes in targets of drugs using ICD10 codes by Genome for Repositioning drugs software.

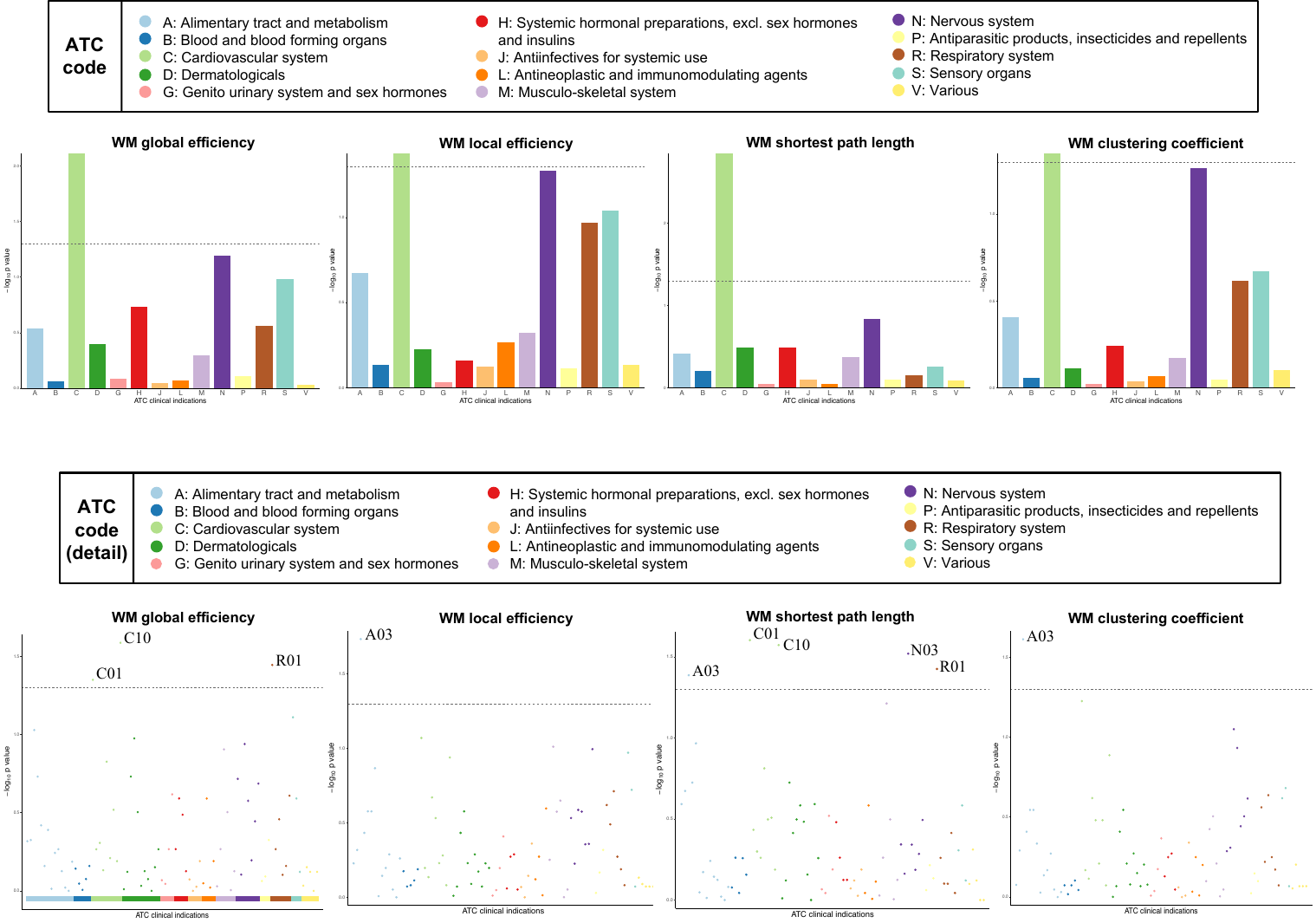

**Supplementary Figure 8.** Enrichment of WM network genes in targets of drugs using ATC codes by Genome for Repositioning drugs software.

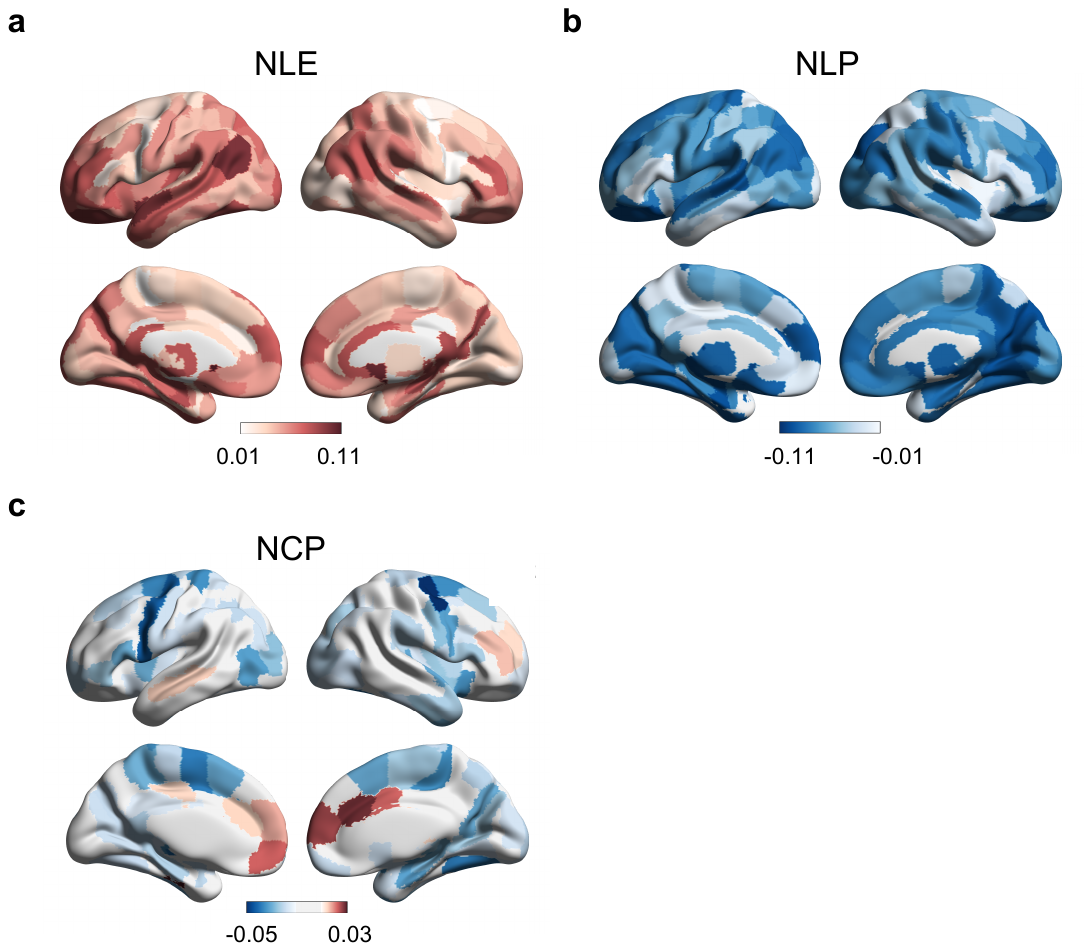

**Supplementary Figure. 9** Phenotypic correlations between intelligence and nodal properties of WM network are illustrated for (a) nodal local efficiency, (b) nodal shortest path length, and (c) nodal clustering coefficient. Regions showing significant genetic correlations with intelligence (false discovery rate corrected P < 0.05) are highlighted to reflect the magnitude of the correlation: positive correlations in red and negative correlations in blue.

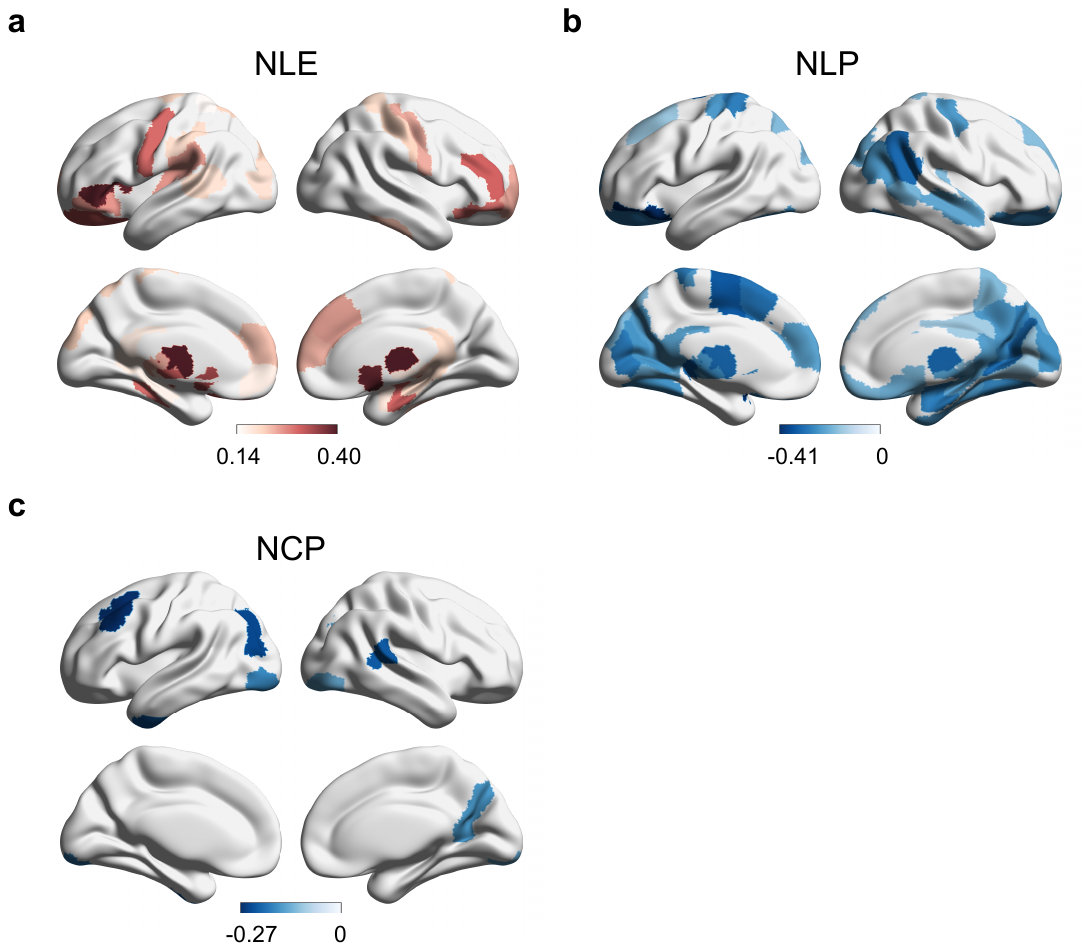

**Supplementary Figure. 10** Genetic correlations between intelligence and nodal properties of WM network are illustrated for (a) nodal local efficiency, (b) nodal shortest path length, and (c) nodal clustering coefficient. The genetic correlations were estimated using LDSC. Regions showing significant genetic correlations with intelligence (P < 0.05) are highlighted to reflect the magnitude of the correlation: positive correlations in red and negative correlations in blue. WM, white matter; LDSC, LD-score regression.

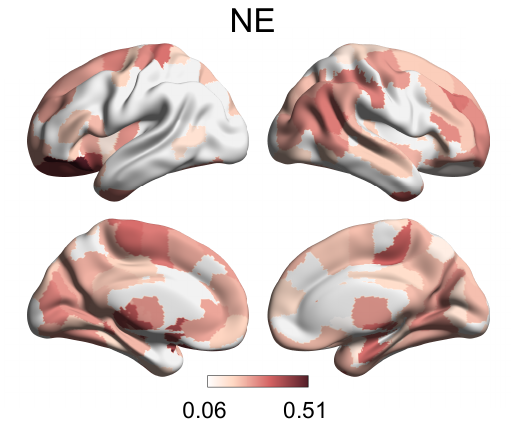

**Supplementary Figure. 11** Genetic correlations between intelligence and nodal efficiency of WM network. The genetic correlations were estimated using LDSC. Regions showing significant genetic correlations with intelligence (P < 0.05) are highlighted. WM, white matter; LDSC, LD-score regression; NE, nodal efficiency.

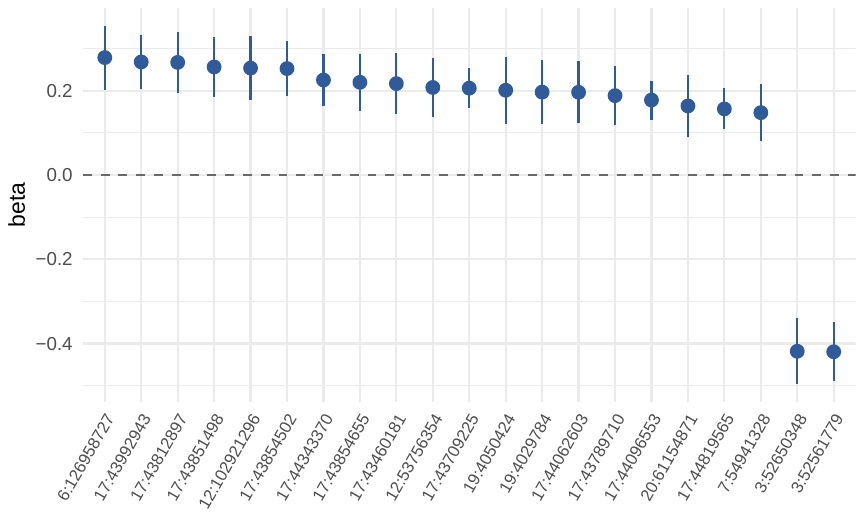

**Supplementary Figure. 12** Causal effects of Eg on intelligence using each of the SNPs after adjusting for multiple testing using FDR.

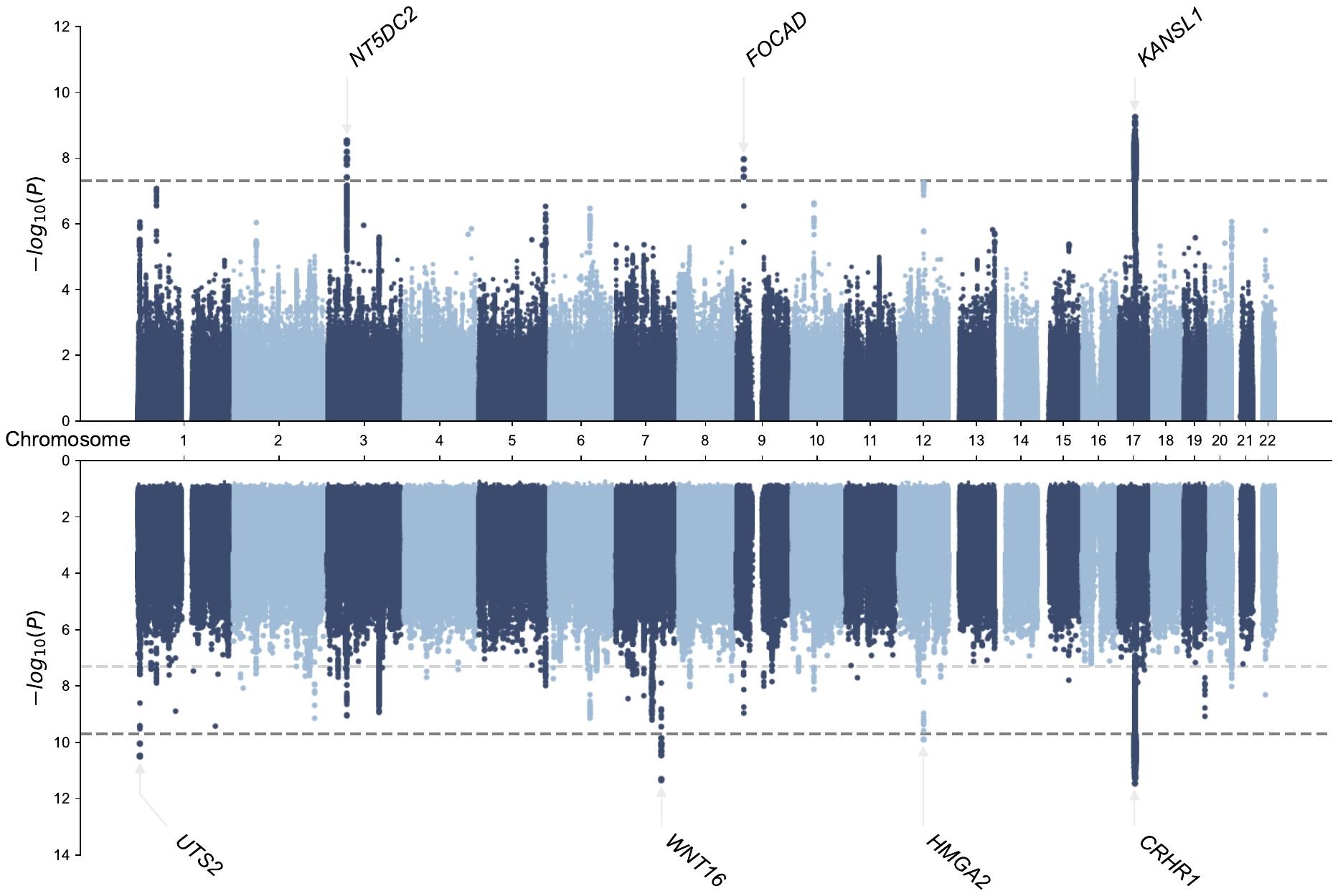

**Supplementary Figure. 13** Manhattan plots with genetic variants identified through univariate GWAS of global efficiency (upper) and multiple univariate GWAS of nodal efficiency across 246 regions with min-P approach (lower) in the first half-sample dataset. The grey lines indicate genome-wide significance threshold (P < 5 × 10–8), and Bonferroni-corrected threshold for the multiple univariate GWAS (P < 2 × 10–10). The independent significant variants are annotated by their nearest genes.

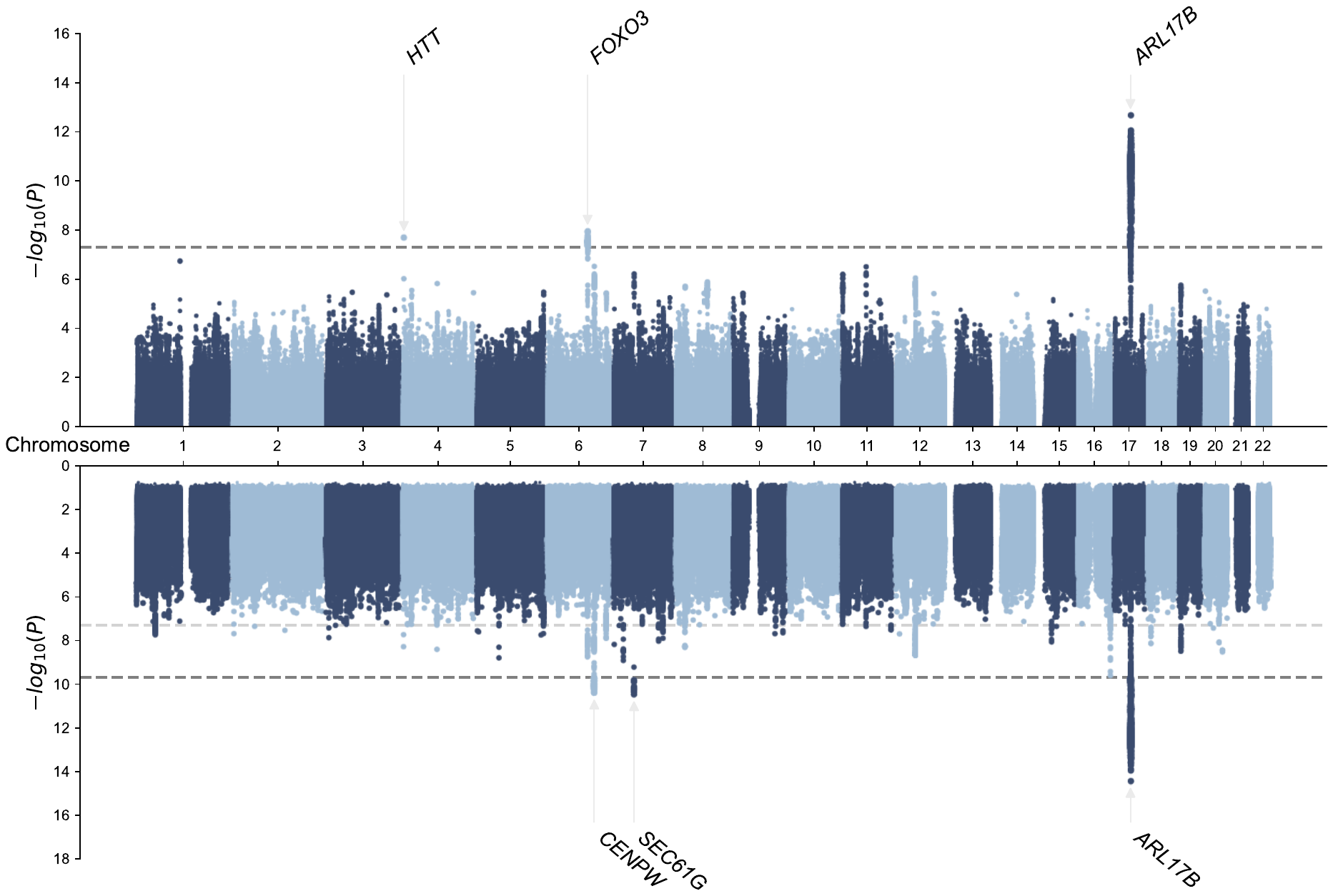

**Supplementary Figure. 14** Manhattan plots with genetic variants identified through univariate GWAS of global efficiency (upper) and multiple univariate GWAS of nodal efficiency across 246 regions with min-P approach (lower) in the second half-sample dataset. The grey lines indicate genome-wide significance threshold (P < 5 × 10–8), and Bonferroni-corrected threshold for the multiple univariate GWAS (P < 2 × 10–10). The independent significant variants are annotated by their nearest genes.

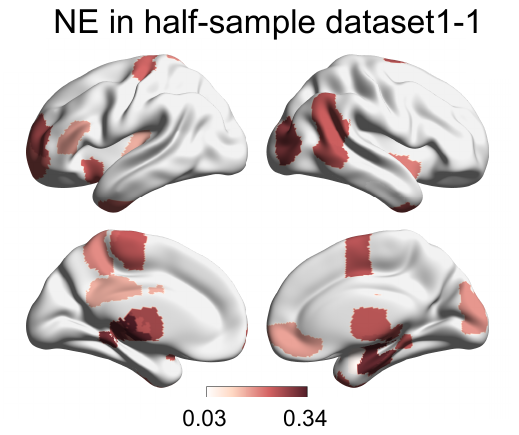

**Supplementary Figure. 15** Genetic correlations between intelligence and nodal efficiency of WM network across 246 regions in the first half-sample dataset. The genetic correlations were estimated using LDSC. Regions showing significant genetic correlations with intelligence (false discovery rate corrected P < 0.05) are highlighted. WM, white matter; LDSC, LD-score regression; NE, nodal efficiency.

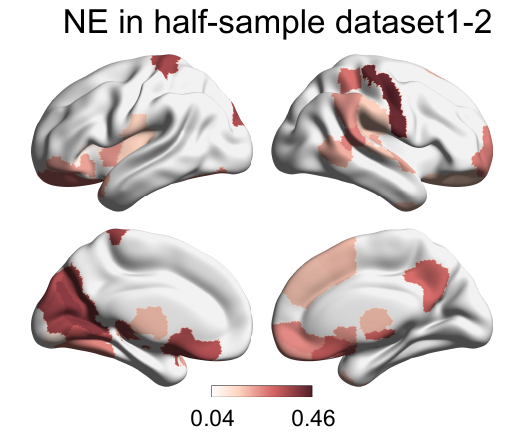

**Supplementary Figure. 16** Genetic correlations between intelligence and nodal efficiency of WM network across 246 regions in the second half-sample dataset. The genetic correlations were estimated using LDSC. Regions showing significant genetic correlations with intelligence (false discovery rate corrected P < 0.05) are highlighted. WM, white matter; LDSC, LD-score regression; NE, nodal efficiency.

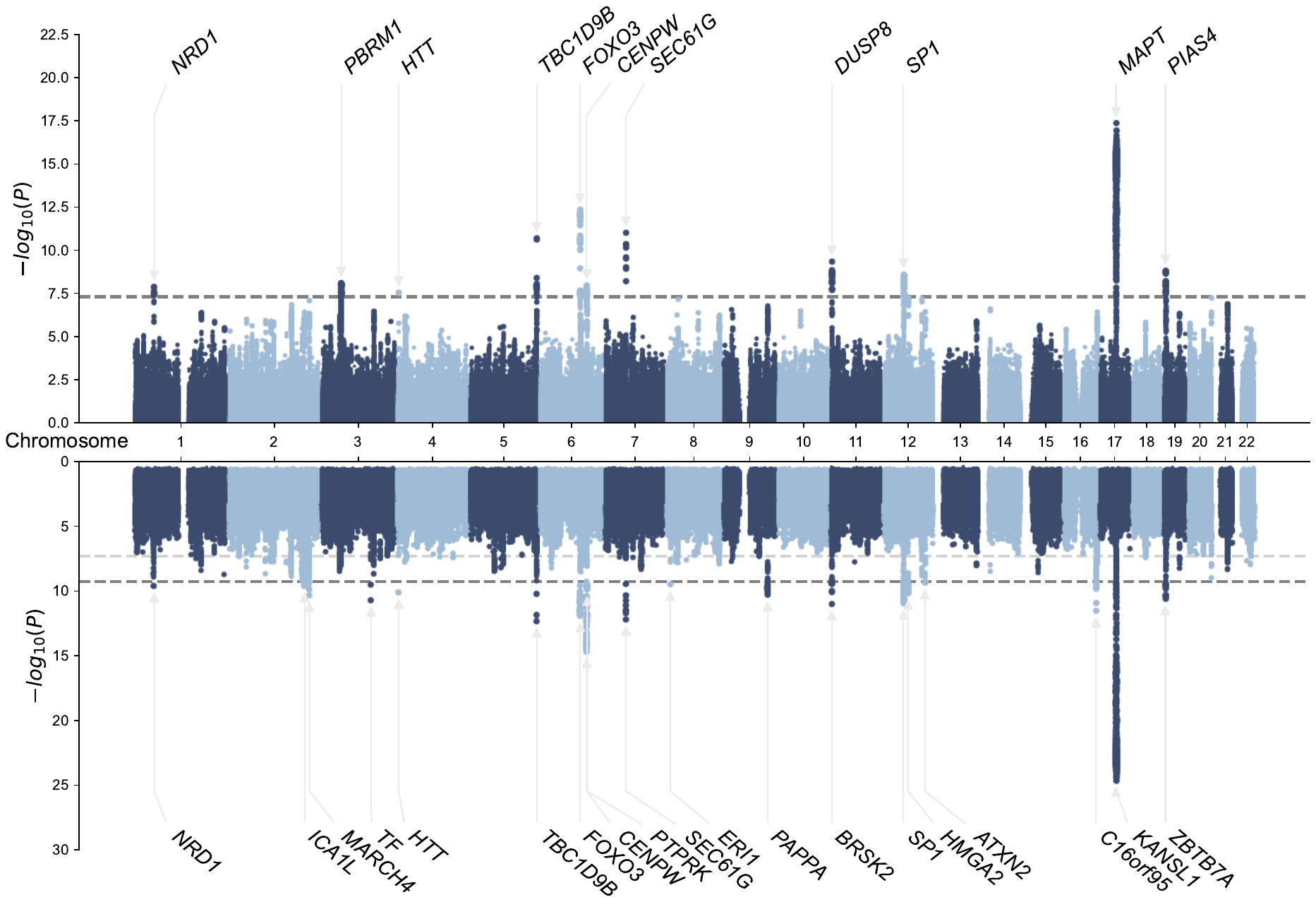

**Supplementary Figure. 17** Manhattan plots with genetic variants identified through univariate GWAS of global efficiency (upper) and multiple univariate GWAS of nodal efficiency across AAL 90 regions with min-P approach (lower). The grey lines indicate genome-wide significance threshold (P < 5 × 10–8), and Bonferroni-corrected threshold for the multiple univariate GWAS (P < 5.6 × 10–10). The independent significant variants are annotated by their nearest genes.

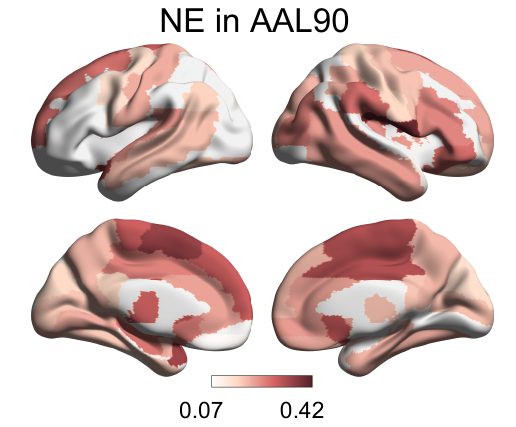

**Supplementary Figure. 18** Genetic correlations between intelligence and nodal efficiency of WM network in AAL90 atlas. The genetic correlations were estimated using LDSC. Regions showing significant genetic correlations with intelligence (false discovery rate corrected P < 0.05) are highlighted. WM, white matter; LDSC, LD-score regression; NE, nodal efficiency.

### References

1.	Alfaro-Almagro, F., et al., *Image processing and Quality Control for the first 10,000 brain imaging datasets from UK Biobank.* Neuroimage, 2018. **166**: p. 400-424.

2.	Jbabdi, S., et al., *Model-based analysis of multishell diffusion MR data for tractography: how to get over fitting problems.* Magn Reson Med, 2012. **68**(6): p. 1846-55.

3.	Fan, L., et al., *The Human Brainnetome Atlas: A New Brain Atlas Based on Connectional Architecture.* Cereb Cortex, 2016. **26**(8): p. 3508-26.

4.	Cook, P., et al. *Camino: open-source diffusion-MRI reconstruction and processing*. in *14th scientific meeting of the international society for magnetic resonance in medicine*. 2006. Seattle WA, USA.

5.	Wang, J., et al., *GRETNA: a graph theoretical network analysis toolbox for imaging connectomics.* Front Hum Neurosci, 2015. **9**: p. 386.

6.	Watts, D.J. and S.H. Strogatz, *Collective dynamics of 'small-world' networks.* Nature, 1998. **393**(6684): p. 440-2.

7.	Maslov, S. and K. Sneppen, *Specificity and stability in topology of protein networks.* Science, 2002. **296**(5569): p. 910-3.

8.	Humphries, M.D. and K. Gurney, *Network 'small-world-ness': a quantitative method for determining canonical network equivalence.* PLoS One, 2008. **3**(4): p. e0002051.
